# Architectural Immunity: ants alter their nest networks to fight epidemics

**DOI:** 10.1101/2024.08.30.610481

**Authors:** Luke Leckie, Mischa Sinha Andon, Katherine Bruce, Nathalie Stroeymeyt

**Affiliations:** School of Biological Sciences, University of Bristol, UK

**Author notes:** Corresponding authors: LL; NS.

## Abstract

In animal groups, spatial heterogeneities shape social contact networks, thereby influencing the transmission of infectious diseases. Active modifications to the spatial environment could thus be a potent tool to mitigate epidemic risk. We tested whether *Lasius niger* ants modify their nest architecture in response to pathogens by introducing control- or pathogen-treated individuals into nest-digging groups, and monitoring three-dimensional nest morphogenesis over time. Pathogen exposure led to an array of architectural changes including faster nest growth, increased spacing between entrances, transmission-inhibitory changes in overall nest network topology, and reduced chamber centrality. Simulations confirmed that these changes reduced disease spread. These results provide evidence for architectural immunity in a social animal and offer insights into how spatial organisation can be leveraged to decrease epidemic susceptibility.

## INTRODUCTION

Animal and human networks influence the spread of disease from local to global scales. For example, the properties of contact and social networks determine the risk and severity of epidemics within social groups^1–3^; urban and metapopulation networks shaped by the layout of buildings, cities and natural habitats influence social interaction patterns and thus affect disease transmission dynamics^4–7^; and global transportation networks facilitate the long-range transmission of disease by connecting distant populations^8^.

There is increasing evidence that modifying social contact networks is an effective intervention strategy against epidemics used in a broad range of animal and human societies. For example, several species including ants, humans, guppies, mice and mandrills avoid infected conspecifics, thereby reducing disease transmission rates^9–11^. Furthermore, ants and humans respond to pathogens by increasing the compartmentalisation of their social networks to limit pathogen spread across the group^2,12^. In addition to modifying their *social* networks, human societies have also used modifications to their *spatial* networks as an active means to reduce disease transmission. For example, the expansion of urban spaces and the separation of cities into functional zones were used as preventative measures against outbreaks of the bubonic plague in the 1300s and cholera in the 1800s, respectively^13^. More recently, active network-based interventions have been proposed which target the spatial properties of transport networks, city layouts, and building architecture to control modern human pandemics such as COVID-19^6,13–15^ Yet, due to the lack of empirical data, it is still unclear which spatial interventions may be most effective at limiting pathogen spread whilst preserving the functioning of society. Animal societies that inhabit complex built structures could provide a unique opportunity to study evolved solutions to this challenge. However, there is as yet no evidence that non-human animals actively modify their spatial surroundings to mitigate epidemic risk, even though such interventions could be highly effective.

Here, we investigate the possible role of architectural changes in nest layout as a disease defence strategy in the black garden ant *Lasius niger*. Nest-building social insects are an ideal system to explore for evolved architectural disease-targeting interventions in animal societies. Indeed, because high interaction rates between related colony members favour the transmission of infectious pathogens, social insects have evolved a large suite of collective mechanisms for disease defence that confer ‘social immunity’, including active modifications to colony social interaction networks^2,16^. Furthermore, built ant nests can demonstrate a high degree of complexity, with specialised chambers housing food, brood, reproductive individuals or waste connected by tunnels into underground networks which successfully isolate potential infectious sources^17,18^. However, whilst previous work has indicated that nest morphogenesis is influenced by the presence of fungal spores in the soil^19^, it is currently unknown whether ants respond to infectious threats by actively modifying their nest architecture to reduce disease transmission.

Here, we test this hypothesis using a combination of pathogen exposure, video recording, X-ray micro-computed tomography (micro-CT), spatial network analysis and agent-based simulations of disease transmission within nests. Our results indicate that pathogen exposure induces ants to excavate nest networks with increased transmission-inhibitory properties, and to place chambers in less central positions. Simulations demonstrate that these interventions reduce disease transmission. This study reveals an unexplored aspect of social immunity and offers new insights into how architectural design might be used for epidemic mitigation.

## RESULTS

To investigate how pathogen exposure influences nest digging by groups of *L. niger* ants, we allowed groups of 180 workers to excavate a new nest in a digging arena. One day after the onset of excavation, we introduced 20 additional workers either exposed to the fungal entomopathogen *Metarhizium brunneum* (pathogen-exposed nests, n=10) or treated with a sham solution (control nests, n=10) into the digging arena. We then monitored excavation for an additional six days using video-recording of surface activity and non-destructive micro CT scans of the internal nest structure (see Fig. 1A-B, Movie S1 and Supplementary Materials).

**Figure 1.**
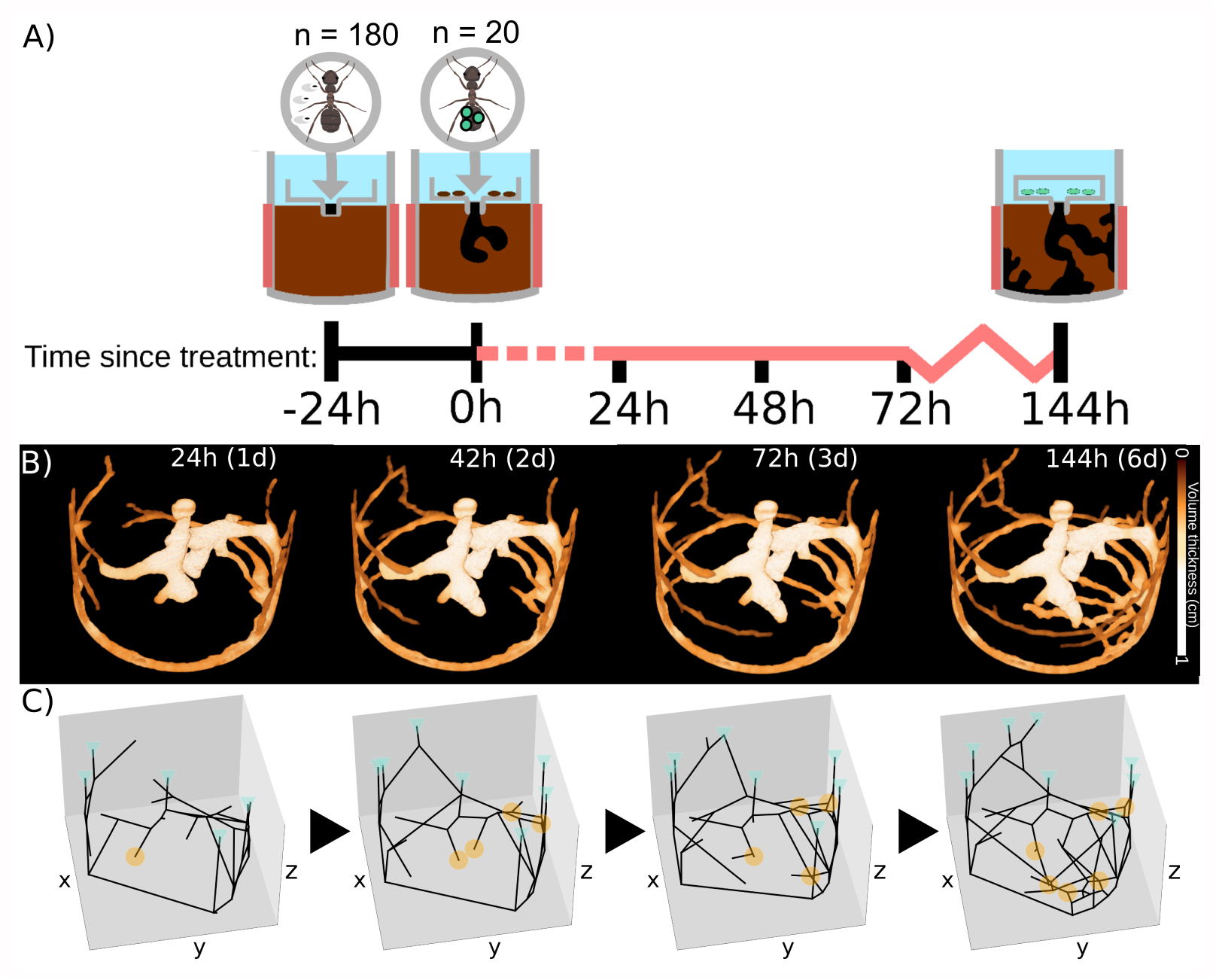
Experimental protocol and nest network extraction. **A:** 180 ants with 25mg early-instar brood and 15 pupae were introduced into a soil-filled container (digging arena) and allowed to excavate a nest freely (-24h). After 24 hours, a CT-scan of the nest was taken (*baseline scan*), then 20 shamor 20 pathogen-treated ants were introduced near the nest entrance (0h). Colonies were allowed to excavate the nest for six additional days and CT-scans were taken 24, 48, 72, and 144 hours (1, 2, 3 and 6 days) after treatment. **B:** CT-scan reconstructions of nest volumes just before treatment (0h), and one day (24h) and 6 days (144h) after treatment. Volume thickness is encoded by increasing colour brightness. **C:** Three-dimensional spatial networks automatically extracted from the nest CT scans. Cyan triangles: nest entrances; orange circles: nest chambers; black lines: tunnels (network edges).

### Pathogen exposure influences surface activity and surface properties of the nest

To investigate the influence of treatment on individual-level activity at the surface, we recorded the number of treated and untreated workers leaving the nest via the main (central) entrance (see Methods). The exit rate of both untreated and treated workers decreased significantly over time in both treatments. However, whilst the exit rate of untreated workers was unaffected by treatment, treated workers exited the main nest entrance at a significantly higher rate in pathogen-exposed than in control nests (Fig. S1; Table S1; linear mixed-effects model (LMM), effect of time, untreated workers: *χ*^2^=208.74, df=1, *p<*0.0001; treated workers: *χ*^2^=6.30, df=1, *p=*0.012; effect of treatment, untreated workers: *χ*^2^=0.021, df=1, p*=*0.89; treated workers: *χ*^2^=6.53, df=1, p*=*0.011; interaction treatment*×*time: *χ*^2^*≤*2.11, p*≥*0.14 in both untreated and treated workers). This indicates that pathogen exposure increases the surface activity of directly-treated workers, but not of their nestmates. Furthermore, we found that entrances were spaced further apart from one another in pathogen-treated than in control nests (Fig. 2, LMM, effect of treatment throughout experiment: *χ*^2^=4.50, df=1, p*=*0.034; Fig. 3A, LMM, effect at 6 days: entrances 0.62±0.30cm further apart in pathogen-treated nests, *χ*^2^=4.19, df=1, p*=*0.041). Increased spacing between entrances could lead to decreased contact rates between individuals at the surface, and has been used as a strategy to mitigate disease transmission in human buildings^20^.

**Figure 2.**
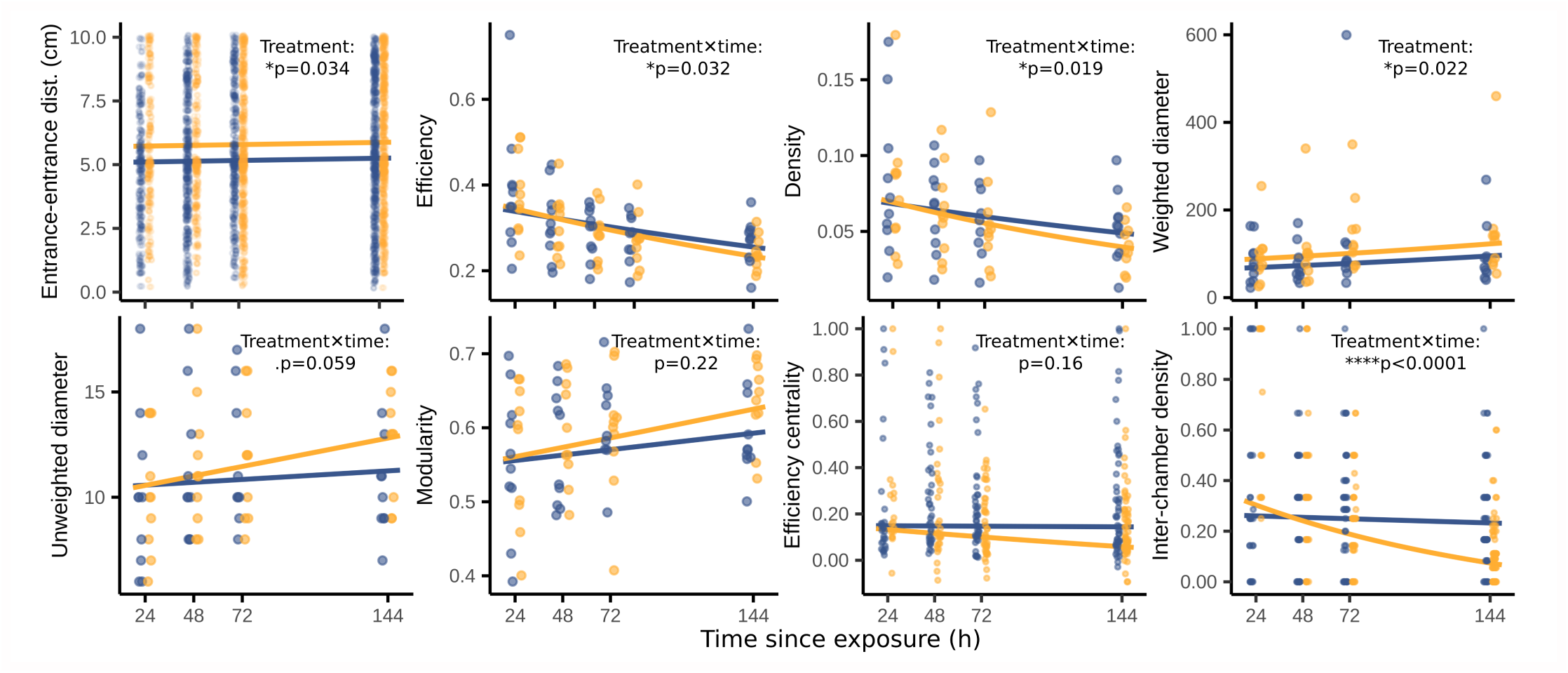
Nest and chamber architectural properties as a function of time. Lines represent LMM fits for control (blue) and pathogen-treated colonies (orange), backtransformed, where appropriate. Significance of the main effect of treatment or the interaction between treatment and time (Treatment*×*time) on a nest property are indicated by p-values. For entrance-entrance distance, each point represents a pairwise measure of euclidean dis(*2*) tance between a new entrance and any other entrance (n=2529 pairs). For efficiency, density, weighted diameter, unweighted diameter and modularity, each point represents one nest (n=79). For efficiency centrality and inter-chamber density, each point represents one chamber (n=336). Efficiency, density, weighted diameter,efficiency centrality and inter-chamber density were log-transformed for statistical analyses. Definitions of all properties are provided in Table 1 and model coefficients and exact p-values in Table S1.

**Table 1.**
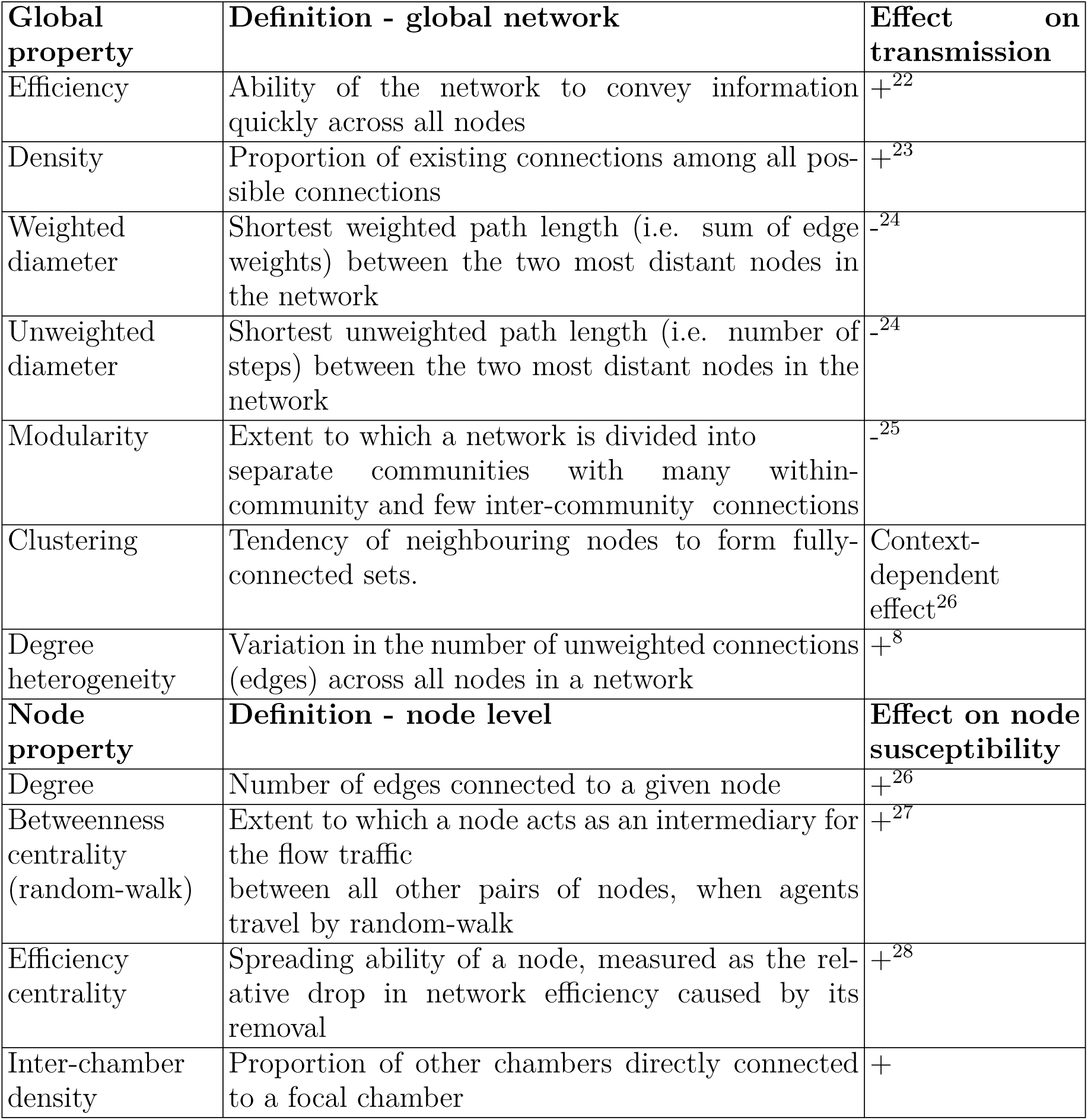
Summary of network properties measured from nest networks and their predicted effects on disease transmission. . + indicates enhancement; - indicates inhibition.

**Figure 3.**
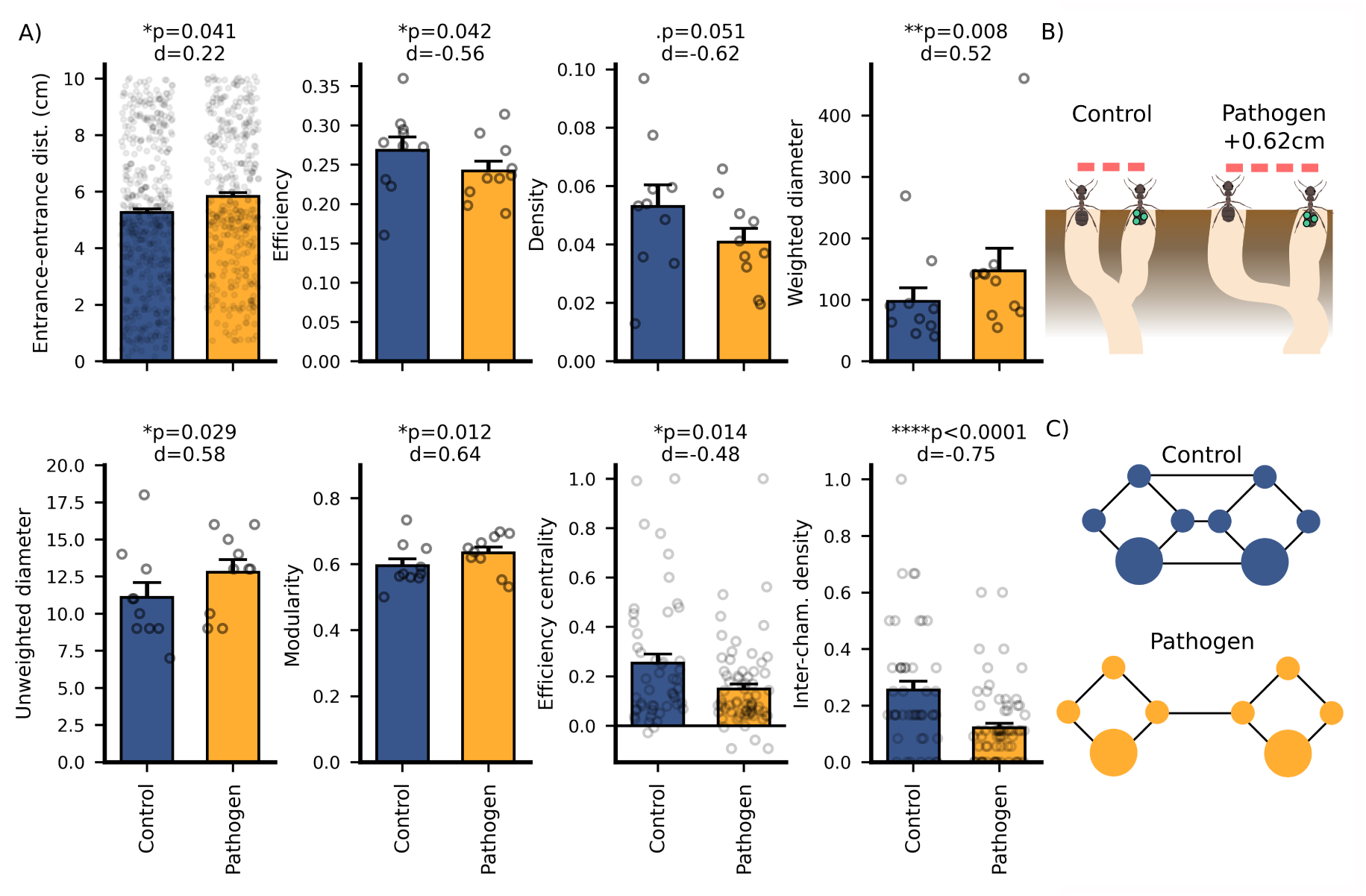
Pathogen-induced architectural modifications 6 days after treatment. **A:** Bars and whiskers show the means and standard errors for each property. Points represent individual data points (entrance-entrance distance (dist.): n=2529 pairs of entrances; efficiency, density, weighted diameter, unweighted diameter, modularity: n=20 nests; efficiency centrality, inter-chamber density: n=336 chambers). Significance of the main effect treatment at day 6 (LMM) and Cohen’s d effect size (|d| 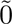.2: weak effect; |d| 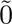.5: medium effect; |d| 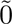.8: strong effect) are indicated above each plot. Model coefficients and exact p-values are provided in Table S1. **B:** Illustration of entrance spacing mediating reduced interaction rate (not to scale). Entrances in pathogen-exposed nests (right) were spaced on average 0.62cm (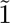 to 1.5 *L. niger* worker body lengths) further apart than in control nests (left), which could reduce the rate of interactions at the surface. **C:** Simplified illustration of between-treatment differences in control (blue) and pathogen-exposed (orange) architectures after 6 days. Small circles represent junctions, large circles represent chambers, and black lines represent tunnels. Pathogen-exposed nests had higher (weighted and unweighted) diameter and modularity, but lower density and network efficiency, and chambers had lower efficiency centrality and inter-chamber density.

### Pathogen exposure influences overall nest growth

To study the influence of pathogen exposure on nest growth, we automatically identified nest chambers, entrances, tunnels, junctions, and dead-ends from the micro-CT scans and measured their volumes (Fig. 1; see Methods and Supplementary Materials for detail). We first compared these and all other measures mentioned below in pre-exposure scans (0h), i.e. *before* treatment was applied. We found no significant differences between treatment groups for any measure, confirming that any differences in post-exposure nests were caused by pathogen exposure rather than due to random initial fluctuations between treatment groups (Table S2).

As higher nest volumes should naturally be associated with lower ant density and thus lower rates of physical contacts between individuals, we predicted that pathogen-exposed groups should excavate nests at a greater rate. In agreement with this prediction, we found that the overall nest volume increased significantly faster in pathogen-exposed than in control nests (Fig. S2, LMM, interaction treatment*×*time: *χ*^2^=5.78, df=1, *p=*0.016). This was not due to differences in the rate of increase of the number and/or volume of nest chambers, or in the rate of increase of the number of nest entrances (Fig. S2; LMM, interaction treatment*×*time: *χ*^2^*≤*1.80, df=1, p*≥*0.18 for all variables). Instead, pathogen exposure led to a significant increase in the rate of tunnel formation (Fig. S2, LMM, interaction treatment*×*time: *χ*^2^=5.49, df=1, *p=*0.019). Although these effects did not lead to detecTable differences in nest volume or tunnel number between treatments by the end of the experiment (Fig. S2, LMM effect of treatment at 6 days, volume: *χ*^2^=0.23, df=1, *p=*0.63; tunnel number: *χ*^2^=0.36, df=1, *p=*0.55), they indicate that pathogen-exposure may lead to changes in the size and connectivity of the nest over longer periods of time.

### Pathogen exposure influences the overall nest network topology

To test whether pathogen-exposed groups of *L. niger* workers alter the topology of their nest network to decrease epidemic risk, we identified properties known to influence disease transmission in social networks^1,2,21^ (Table 1), and measured them in the spatial nest networks consisting of chambers, junctions, dead-ends and entrances (nodes) connected by tunnels (edges; Fig. 1C). We predicted that pathogen-exposed nest networks would display increased transmission-inhibitory properties and decreased transmission-enhancing properties compared to control nests.

In agreement with this prediction, we found that nest network efficiency and density (transmission-enhancing properties) decreased at a significantly higher rate in pathogen-exposed than in control nests (Fig. 2, LMM, effect of treatment*×*time, efficiency: *χ*^2^=4.58, df=1, p*=*0.032; density: *χ*^2^=5.52, df=1, *p=*0.019). Both properties were reduced in pathogen-exposed nests by 6 days after treatment, though this was marginally non-significant for density (Fig. 3A, LMM, effect of treatment at 6 days, efficiency: *χ*^2^=4.125, df=1, p*=*0.042; density: *χ*^2^=3.82, df=1, p*=*0.051). In addition, we found that the weighted diameter of the nest network (transmission-inhibitory property) was significantly higher in pathogen-exposed than in control nests throughout the experiment (Fig. 2, LMM, effect of treatment throughout experiment: *χ*^2^=5.26, df=1, p*=*0.022; Fig. 3A, LMM, effect at 6 days: *χ*^2^=6.97, df=1, p*=*0.008). Furthermore, the unweighted diameter and modularity of nest networks (transmission-inhibitory properties) tended to increase faster in pathogen-exposed than in control nests, and both properties were significantly higher in pathogen-exposed nests by the end of the experiment (Fig. 2, LMM, effect of treatment*×*time, unweighted diameter: *χ*^2^=3.57, df=1, *p=*0.059; modularity: *χ*^2^=1.48, df=1, *p=*0.22; Fig. 3A, LMM, effect of treatment at 6 days, unweighted diameter: *χ*^2^=4.79, df=1, *p=*0.029; modularity: *χ*^2^=6.34, df=1, *p=*0.012). No other transmission-relevant properties of the overall nest network were affected by treatment (Table S1; LMM for clustering and degree heterogeneity: effect of treatment*×*time, *χ*^2^*≤*0.07, df=1, p*≥*0.51; effect of treatment at 6 days: *χ*^2^*≤*1.35, df=1, p*≥*0.25 in all tests). Overall, these results indicate that pathogen exposure leads to multiple changes in the overall topology of the nest networks which should decrease pathogen transmission.

### Pathogen exposure influences the topological position of nest chambers

Previous studies of transport and social networks have shown that highly-populated nodes and nodes with high network centrality (e.g. *degree*, *betweenness* and *efficiency centrality*, see Table 1) are both at high risk of contamination and highly influential on onward spread to the rest of the network^2,29–31^. As nest chambers contain a large portion of the nest population including valuable and vulnerable colony members such as the queen, young adults and the brood, we predicted that pathogen-exposed groups would build chambers in positions associated with lower exposure risk and lower spreading ability, characterised by lower network centrality and reduced connections to other chambers.

Most chambers had a degree of three, and neither chamber degree nor betweenness differed between treatments (Table S1, Poisson GLMM (degree) and LMM (betweenness), interaction time*×*treatment and main treatment effects: *χ*^2^*≤*0.0053, df =1, p*≥*0.95 in all tests; Poisson GLMM (degree) and LMM (betweenness) at 6 days: *χ*^2^*≤* 0.20, df =1, p*≥*0.66). However, in agreement with our prediction, the efficiency centrality of nest chambers tended to decrease faster in pathogen-exposed than in control nests and was significantly reduced in pathogen-exposed nests by the end of the experiment (Fig. 2, LMM, interaction treatment*×*time, *χ*^2^=1.97, df=1, *p=*0.16; Fig 3A, LMM, effect of treatment at 6 days: *χ*^2^=6.07, df=1, *p=*0.014). Furthermore, the density of connection of chambers to other chambers decreased significantly faster in pathogen-exposed than in control nests, and was significantly reduced in pathogen-exposed nests by 6 days after exposure (Fig. 2, LMM, interaction treatment*×*time: *χ*^2^=25.90, df=1, *p<*0.0001; Fig 3A, LMM, effect of treatment at 6 days: *χ*^2^=16.02, df=1, *p<*0.0001). Altogether, these results indicate that pathogen-exposed ants excavate chambers in positions which should reduce the severity of epidemics.

### Pathogen-induced changes in nest architecture decrease disease transmission

Our results so far indicate that pathogen exposure triggers a range of transmission-inhibitory changes in nest architecture effective by 6 days after treatment (Fig. 3). To formally test whether these changes do decrease the risk of epidemics, we developed an agent-based model which simulated the transmission of an infectious pathogen within the observed nest networks. This model was inspired from previous models of disease transmission in ant nests^2^ and ant traffic in confined space**^chang_nest_^**^2021, 32^(see Methods and Supplementary Materials for detail). In the model, agents moved through the nest and aggregated locally inside nest chambers^18,33^. Pathogen-treated agents initially entered the nest through a randomly-selected entrance, and pathogen transmission occurred as a stochastic process between pairs of agents sharing the same within-nest location (chambers, tunnels, dead-ends and junctions). We simulated the transmission of *M. brunneum* over the experimental nest networks extracted six days after treatment, and found that untreated agents had a significantly lower simulated fungal load and a significantly reduced prevalence of high (more lethal) loads in simulations over pathogen-exposed than control nests (Fig. 4A-B, LMM interaction treatment*×*time, median load: *χ*^2^=49.73, df=1, *p<*0.0001; prevalence of high load *χ*^2^=30.57, df=1, *p<*0.0001).

**Figure 4.**
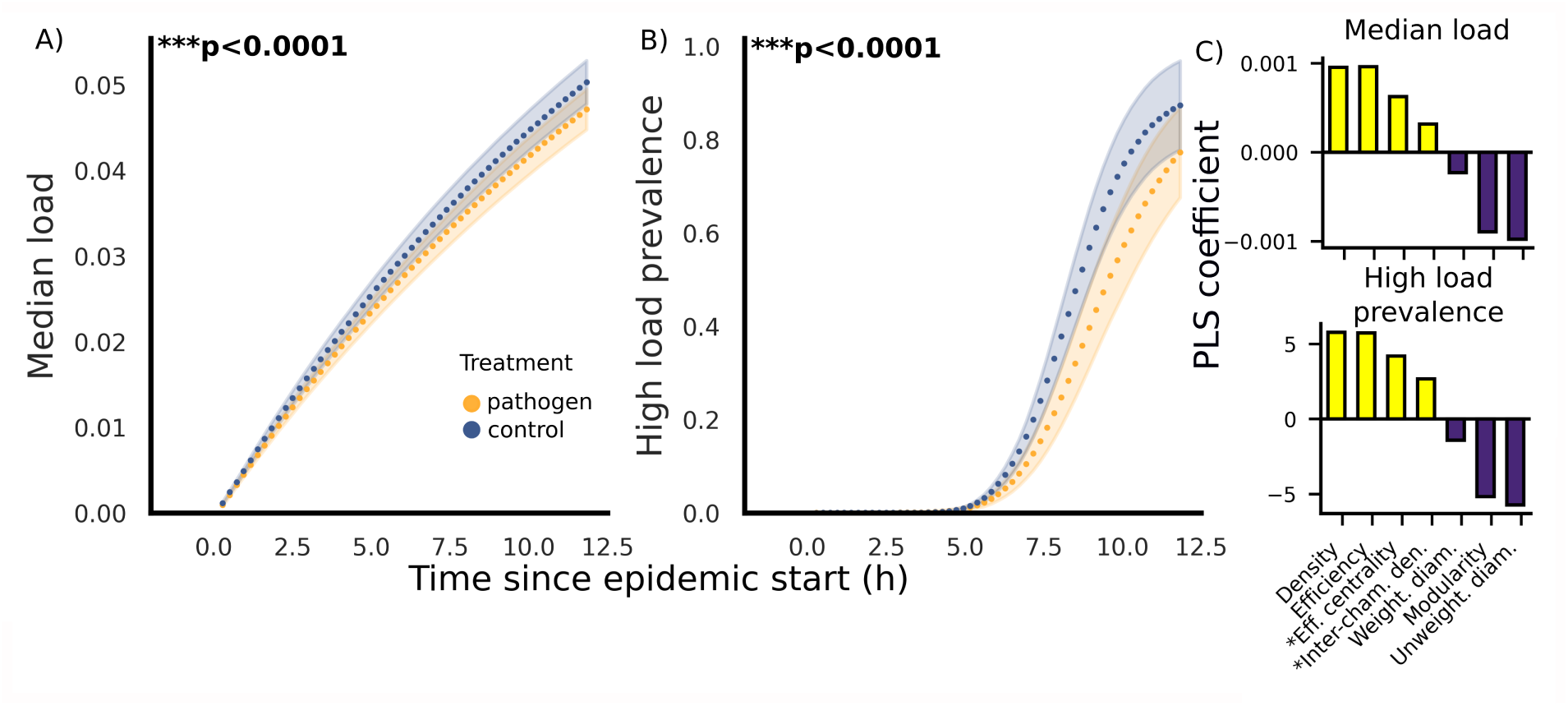
Agent-based simulation results of *M. brunneum* transmission within 6-day nest networks. Median load (**A**) and high load prevalence (**B**) across *untreated* agents in simulations over control (blue, n=10) and pathogen-exposed (orange, n=10) nest networks. 100 simulations were performed on each nest and cross-simulation means were calculated for each nest every 800 seconds after introducing treated agents (n=53 time points), resulting in one data point per nest per time point. Points and shaded areas represent the grand means and standard errors calculated across the 10 nests per treatment at every time point. Details of statistical analyses are provided in Table S1. **C:** bars indicate coefficients from partial least-squares regression (PLS) analysing the relative effects of nest network properties targeted by architectural modifications on simulated median load (top) and high load prevalence (bottom). Positive (resp. negative) values indicate a positive (resp. negative) association between a property and transmission, and bar height is proportional to its relative influence on transmission. Transmission-enhancing properties are indicated in yellow and transmission-inhibitory properties in purple. Chamber properties are highlighted by an asterisk; all other properties apply to the global topology of the nest. All pathogen-induced changes were transmission-inhibitory.

To further tease apart the relative importance of different pathogen-induced architectural modifications in reducing transmission, we applied partial least square regression to the outcome of simulations over all nests. All properties were found to have the expected effects on transmission: presumed transmission-inhibitory properties (weighted diameter, unweighted diameter, modularity and inter-chamber density) were negatively associated with transmission, whilst presumed transmission-enhancing properties (density, efficiency and efficiency centrality) were positively associated with transmission (Fig. 4C). Efficiency, density, unweighted diameter, and modularity (whole-network properties) had the strongest associations with transmission; chamber properties had weaker associations, with efficiency centrality showing a stronger effect on transmission than inter-chamber density; and weighted nest diameter had the weakest association with transmission (Fig. 4C).

## DISCUSSION

Group-living animals have evolved collective responses to infectious pathogens that decrease the risk of epidemics^2,9–12^. Most responses reported so far involve changes in social interactions between individuals, ranging from social care to self-isolation, social distancing, and social network reorganisation^34^. Our results indicate that these social changes are complemented by spatial modifications to the environment, which had previously only been shown in humans.

Our approach, combining serial micro-CT scanning and spatial network analysis, revealed three types of responses by groups of *Lasius niger* ants challenged with the fungal pathogen *M. brunneum* in the early stages of nest digging. First, pathogen-exposed workers increased their surface activity, which likely reflects self-isolation and social distancing^2,11,35^. Second, pathogen exposure led to changes in the geometry of the nest: pathogen-exposed groups increased the spacing of their nest entrances, which should decrease the rate of surface interactions and hence the risk of pathogen transmission; they dug at a faster rate, which should decrease overall nest population density and hence the rate of below-ground interactions; and they excavated more tunnels, which may provide redundant or alternative routes of transport avoiding transmission hubs such as nest chambers^35^. Interestingly, these changes echo architectural and urban interventions in human societies, which have used increased entrance spacing and urban expansion to limit COVID-19 and bubonic plague transmission, respectively^15,20^. Third, pathogen exposure led to changes in the topology of the nest network: pathogen-exposed nests showed increased transmission-inhibitory properties (modularity and diameter) and decreased transmission-enhancing properties (density and efficiency) relative to the control, which should limit disease spread. Furthermore, nest chambers occupied topological network positions associated with lower exposure risk and spreading ability^28^ (lower efficiency centrality and inter-chamber density), which should help contain epidemics and protect high-value and/or vulnerable individuals housed in chambers, such as the queen, young nurses and the brood. Notably, the network properties that the ants alter have previously been predicted to exert a critical influence on epidemic susceptibility in a broad range of animal and human social and spatial networks^21,25^.

Our results contrast with a previous study in the ant *Myrmica rubra*, which detected neither avoidance nor increased digging rate in *Metarhizium*-contaminated soils^19^. However, in this previous study, only the first 40 hours of nest excavation were recorded, which may have been too little to detect clear pathogen-induced effects on digging activity. Furthermore, the authors did find that nests excavated in contaminated soils were more anisotropic, with more/longer tunnels and a smaller main chamber. Although the consequence of these changes for transmission was not considered in that study, the increased nest spatial heterogeneity is likely to limit disease spread^5,36^, suggesting that transmission-inhibitory architectural responses to pathogens may be widespread in ant species.

Simulations of disease transmission over control and pathogen-exposed nest networks confirmed that the pathogen-induced changes in nest architecture successfully inhibit disease transmission within the nest, and hence protect the colony against epidemics. Importantly, these beneficial effects emerged without requiring any assumptions on social organisation or pathogen-induced changes in individual behaviour, showing that nest architecture alone is sufficient to reduce epidemic risk. However, in natural conditions, division of labour, self-isolation, and social distancing are likely to act in synergy with nest architecture to provide enhanced protection against pathogen threats, as they play a key role in *L. niger*’s defences against disease even in simple, single-chamber nests^2,37^.

Most pathogen-induced changes in architecture increased over time, which could be linked with the nest’s development. At the beginning of the experiment, nests were small, which may have limited the scope for implementing architectural changes. At this early stage, self-isolation by pathogen-exposed individuals may therefore be the most effective strategy to prevent the introduction of infectious material into the nest. As nests grow larger, with more chambers, junctions and tunnels, they may become more amenable to topological manipulation, making architectural modification a more potent - and durable - epidemic defence strategy. Indeed, the number of possible network configurations scales non-linearly with the number of nodes^38^. By emphasising social distancing early on and architectural modifications later in nest development, pathogen-exposed ants may dynamically adapt their social immunity to implement the most effective defence.

The mechanisms driving the pathogen-induced changes in nest architecture remain to be established. Although infection may affect digging activity in diseased workers, it is unlikely to be the main cause of the architectural modifications reported here for three reasons: first, changes to entrance-entrance distance and weighted diameter were detecTable as early as 24 hours after exposure (i.e. before the fungus can cause infection^39^); second, nests grew faster in pathogen-exposed than control groups, ruling out a reduction in digging activity as a cause for the changes; and third, pathogen-treated workers spent more time near the surface and should thus have less influence on belowground nest morphology. Instead, the topological changes likely reflect an active shift in decision-making amongst untreated workers about where to excavate entrances, chambers and tunnels. Identifying the fine-grained individual-level mechanisms that lead to this shift in collective decision-making represents an exciting area of future research.

Overall, our results indicate that ants use transmission-inhibitory modifications to their nest architecture as a strategy to reduce epidemic risk upon acute pathogen threat, providing a form of ‘architectural immunity’. These findings could have important implications beyond animal societies, as the architectural changes highlighted here have been tuned for effectiveness over long evolutionary time and could serve as a proof-of-concept or source of inspiration for human disease interventions^2,40^.

## METHODS

### Collection and rearing of colonies

Workers and brood from six mature field colonies (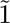1000-2000 workers) of *L. niger* were collected from Bristol, UK between August and October 2022. Colonies were maintained in the laboratory in 41cm (L) x 26cm (W) x 15cm (D) fluoned, soil-filled, containers at 25°C, 65% humidity, with a 12hr day/night cycle. Colonies were fed with ad-libitum protein (*Drosophila hydeii* and cockroaches) and 10% sugar water until the start of the experiment.

### Experimental design

The experiment followed a matched-pairs design: every week, we ran one experimental replicate involving two 200-ant subsets from the same colony, one of which was subjected to the pathogen-exposure treatment and the other to the control treatment. For each replicate, two 200-ant subsets were randomly sampled from the same colony, alongside 25mg early-instar brood and 15 pupae each. 20 random ‘challenge’ individuals were then randomly selected from each subset, marked with a dot of Pactra™ acrylic paint on their gaster, and transferred into separate boxes. The remaining subsets (180 workers each) were kept in 17.5cm (length) x 12cm (width) x 1.5cm (depth) boxes coated with Fluon® to prevent the ants from escaping. Three days after subsetting, each subset of 180 workers was introduced into a digging arena consisting of a 11cm x 18cm (depth) plastic cylinder with fluoned wall and a moistened plaster basis filled with a mixture of sand and clay (10% clay) at 15% hydration, which has been shown to stimulate branched digging in *L. niger* ^41^. The cylinder walls were covered with red filter paper to maintain darkness below the substrate surface. Each cylinder was surmounted by an entrance arena positioned on top of the substrate and consisting of a closed Petri dish (90mm) with a central hole connected to a downward tube (11mm length, 6mm) inserted into the top 9.5mm of the substrate. This ensured that digging was always initiated at the same location (Fig. 1A). A 1.5mm gap between the substrate surface and the Petri dish floor and a 1cm perimeter gap between the Petri dish and the cylinder walls allowed the formation of nest entrances under and around the Petri dish. Subsets were introduced into the entrance arena, then allowed to excavate a nest freely for 24 hours. Baseline architecture was then recorded by taking a non-destructive micro-CT scan of each nest (0 hours post-treatment, Fig. 1A-B; see Methods and Supplementary Materials for detail on CT scanning). Immediately after scanning, the 20 ‘challenge’ individuals associated with each subset were exposed to a live *M. brunneum* spore suspension (‘pathogen-exposed nests’) or to the solvent only (‘control nests’), then introduced inside the entrance arena (see Methods and Supplementary Materials for detail on treatment). Colonies were then allowed to excavate nests for a further 6 days. We recorded post-treatment nest architecture with further CT scans at 24, 48, 72, and 144-hours (i.e. 1, 2, 3 and 6 days) after treatment (Fig. 1A-B). Furthermore, to test the influence of treatment on individual activity, we took five-minute video recordings of the main entrance every hour for the first two hours after treatment, then every two hours thereafter.

A total of 10 experimental replicates (10 pathogen-exposed nests and 10 control nests) were carried out over a period of 10 weeks, which resulted in 100 total CT-scans. Due to a scanning error, we discarded one of these scans from our empirical analyses (the first control replicate at 72 hours). Due to constraints on colony sizes, four colonies were used as sources for two replicates each and two colonies were used as sources for one replicate each; this was accounted for using random effects in all statistical analyses (see below).

### Video analysis

Videos were recorded on a raspberry-Pi IR-CUT infrared camera and analysed by two blinded observers, who counted individuals exiting the initial entrance within each five-minute video recording for the first 53 hours after treatment (n=446 5-minutes clip corresponding to 37h10min of footage in total); after this point, the entrance was frequently obscured by substrate build-up, which prevented further activity monitoring. Treated ant activity was discerned from untreated nestmate activity using the individual paint marks on treated ants. Inter-observer agreement was measured at 100% over five-minute samples taken across a 24-hour sample of test footage.

### Nest network extraction

Micro-CT scan outputs were first transformed into skeletons, which structurally represent the medial lines of the scanned volumes. We then automatically extracted networks from skeletons using a custom Python script (see Supplementary Materials, Figs. S3-4). In these networks, edges were tunnels and nodes were junctions, deadends, entrances and chambers. Edge weights, *w*, between connecting nodes *i* and *j* were given as 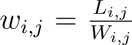, where *L* is tunnel length and *W* is tunnel width. Weights reflect the fact that shorter and wider tunnels should facilitate traffic flow; the higher the edge weight, the more distant (i.e. the less well-connected) the two connected nodes. Node type (entrance, junction, endpoint and chamber) was identified using a semi-automatic pipeline detailed in the Supplementary Materials. In short, chambers were identified first using a combination of manual assignment by blinded observers and algorithmic criteria (enlarged width compared to the surrounding area and with a horizontal base^42^); entrances were identified second based on their own and neighbouring nodes’ elevation; and all remaining nodes were labelled as junctions if they had more than one connection or endpoints if they had a single connection. Finally, nodes were assigned unique identities which were maintained between nest networks at successive time points to allow us to analyse the change in node properties over time.

### Entrance distribution

We determined the spacing between entrances by calculating pairwise Euclidean (i.e. beeline) distances between unique pairs of nest entrances.

### Nest network analysis

All network analyses were performed using Python package NetworkX 2.8.7^43^. For betweenness and modularity, where weight must indicate the strength of connection, edge weights were inverted. Analyses were performed on the largest connected component, in which the majority of tunnels were located (proportion of edges in main component, control: 0.97±0.03; pathogen 0.97±0.05). There was no significant difference between treatment groups in the number of connected components or the proportion of edges in the main connected component, and no chambers were ever located in secondary components (GLMM with poisson distribution, interaction treatment*×*time, number of connected components: *χ*^2^=0.07, df=1, *p=*0.79; LMM interaction treatment*×*time, proportion of edges in main connected component: *χ*^2^=1.60, df=1, *p=*0.21).

To examine whether groups of pathogen-exposed ants modify the global nest topology we measured network efficiency, density, modularity, (unweighted and weighted) diameter, clustering and degree heterogeneity ; for definitions see Table 1. Modularity was measured using the Louvain algorithm^44^.

To examine whether groups of pathogen-exposed ants build chambers in positions with lower spreading ability and lower exposure risk, we used three measures of network node centrality, which are epidemiologically relevant in human spatial networks: degree, random-walk betweenness (hereafter betweenness) and efficiency centrality^14,28^ (see Table 1 for definitions). To account for network size, betweenness and efficiency centrality were normalised by dividing the measured value by the maximum value across all nodes in the focal network.

### Simulation model

#### Model input

We developed a model to simulate the transmission of disease as agents move over the experimental nest networks extracted from CT-scans 6 days after treatment (see Supplementary Materials and Table S3). Nest components possessed spatial properties that influenced agent movement and interactions: tunnels were characterised by their length and width, and chambers by their diameters. Since we were unable to obtain diameter measurements for all chambers, for simulation purposes we used the median chamber diameter of each nest as the diameter of all chambers in that nest.

#### Model initialization

At the beginning of each simulation, we calculated the proportion of the total nest volume occupied by chambers. The same proportion of agents were then distributed equally across the chambers in the focal nest at t=0. The remaining agents were each allocated to a random entrance.

#### Agent movement

Each time step represented one second. Agents moved according to a randomwalk^45^. At each time step, we first considered the movement of agents from nodes into their connecting tunnels. Exiting a node was modelled as a stochastic process with a probability *P_L_* depending on the agent’s location. Agents in junctions, dead-ends and nest-entrances had an exit probability of 1, unless all connecting tunnels were full (see below), in which case their exit probability was zero. For chambers, the exit probability, *P_L_* was given by: 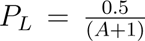, where A is the number of agents in the node^33^. If an agent was determined to exit its current node, it entered a randomly selected tunnel connected to that node. To reflect outside movement, agents leaving entrances randomly selected an entrance to enter, from which they entered a random tunnel. If an agent left the initial entrance, which was enclosed, they re-entered a random tunnel connecting to the entrance.

Agents moved through tunnels and between entrances at half a body length per second (0.2cm/s)^46^. For simplicity, we consider tunnels to be two-dimensional^32,33^ and to have a maximum density of 1.33 agents/cm^2^, which we used to determine the agent capacity of a tunnel^66^. Hence, after entering a tunnel or leaving the nest agents remained in the tunnel or ‘outside’ for a total duration determined by 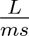, where *L* is the length of the tunnel or the euclidean distance between entrances and *ms* is agent movement speed, before moving into the subsequent node. If the agent re-entered the same entrance, *L* was randomly drawn from a uniform distribution between 0 and 9, which bound the possible euclidean distances required to travel to, and return from, a random point along the radius of the entrance arena, from the initial nest entrance. If the subsequent node was a junction and it was already occupied by two agents, agents had an equal probability of either waiting for one timestep or turning around. Agents that turned around spent an additional 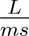, timesteps within that tunnel, before attempting to re-enter the node from which they originally entered the tunnel.

The initial 180 agents were first allowed to move around the nest for 21600 timesteps (6 hours) in the absence of disease. At this point, 20 *treated* agents were added to the simulation. *Treated* agents were initialised with a spore load of 1 and each was allocated to a randomly selected nest entrance. *Treated* agents then successively entered the tunnel connected to their allocated entrance, with a delay of 2 seconds between successive agents (time for an agent to move one body length). After the introduction of treated ants, spore transmission between pairs of agents was allowed to occur within the nest.

#### Contact and spore transmission

Within each below-ground component (chambers, junctions, dead-ends and tunnels), interactions were considered sequentially across randomly shuffled agent pairs. At each time step, the number of pairwise interactions *N_i_* occurring within tunnels and chambers that contained at least two agents was randomly drawn from a Poisson distribution with parameter *\* A*, where is the mean number of interactions per individual and per second (0.0208.ant^-1^.s^-1^), and *A* is the number of agents within the focal component. We parameterized using unpublished automated tracking data from 30 *L. niger* colonies, ranging between 100 and 180 ants in size. We then repeatedly drew random pairs of agents from the focal component and labelled them as interacting until *N_i_* was reached or until each agent was involved in a maximum of one interaction each. Within junctions *N_i_* was always one. Transmission between pairs of interacting agents was then modelled as a stochastic event using the equations and parameter values developed in a previous model of transmission of *M. brunneum* in *L. niger* ants^2^. In short, transmission could occur with probability (0.34) between pairs involving at least one agent with a load above the infectiousness threshold (0.00024). If transfer did occur, an amount of spores *S* was transferred unidirectionally from the agent with the higher load to the agent with the lower load, calculated according to the following formula: *S* = Δ*λ ∗ dt ∗ γ*, where Δ*λ* is the difference in load, *γ* is the transmission rate (0.000345), and *dt* is the duration of the contact (1 timestep).

The model was allowed to run with transmission interactions for an additional 43200 timesteps (i.e. 12 hours) after the introduction of *treated* agents. We limited our simulation time to 12 hours since our experiment determined nests to change considerably from day to day, limiting the timeframe over which static nest representations are relevant. All individual spore loads were recorded every 800 timesteps. At each recorded time step, we determined the median simulated spore load across all untreated agents as well as the ‘high load prevalence’, i.e., the proportion of untreated agents carrying a high, potentially lethal load (as parameterised in^2^). We performed 100 simulations for each nest (control: n=10; 144h, pathogen-exposed: n=10) and calculated a crosssimulation mean of the two variables of interest (median untreated load and high load prevalence) for each nest at each recorded timestep, to be used in onward statistical analyses.

### Statistical analyses

All statistical analyses were performed using R version 4.3.0 and used two-sided hypothesis tests. Unless otherwise specified, all mixed-effect models listed below included source colony and experimental replicate as random factors.

We compared treatment-induced changes in surface activity, nest architecture and simulated transmission using linear mixed-effects models (LMM; continuous data) or generalised linear mixed-effects models (GLMM) with Poisson distribution (count data) using the R package *lme4*. To ensure results were not influenced by initial differences, we first compared pre-treatment networks extracted from the 0h-scans using simple models with treatment as the only main effect (Table S2). To investigate the influence of treatment on nest architecture over time, we fitted models from 24-hours to 144-hours post-exposure including time (continuous variable), treatment and their interaction as main effects, and subset identity and - for models of chamber properties - chamber identity as additional random factors. To investigate the influence of treatment on nest architectures at the end of the experiment, we fitted models on nest properties extracted six days after treatment, with treatment included as the only main effect. Additionally, at this time-point we used Cohen’s d to measure the effect size of treatment-induced changes. Cohen’s d was calculated as the mean over pathogen-treated nests minus the mean over control nests divided by the pooled standard deviation (|d| 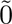.2: weak effect; |d| 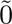.5: medium effect; |d| 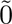.8: strong effect)^47^. Models of surface activity were fitted separately for treated and untreated workers. For the simulated transmission data on median agent load and high load prevalence, we used LMMs that included median load as a response and treatment and timestep as interacting main effects.

For all LMM, diagnostic plots (residuals vs. fitted values and QQ plots) and Shapiro-Wilk tests were used to check whether test assumptions of normality and homogeneity of variances were met. Data were transformed wherever necessary to ensure normal distribution of residuals. Hypothesis testing was performed using type III ANOVA with Wald’s test on the fitted models.

To analyse the relative contribution of nest properties in modulating transmission, we used Python package scikit-learn to fit partial least square regressions on the simulation outcome (median untreated load and high load prevalence) obtained from all 6-day nests as a function of the seven belowground nest properties found to differ between pathogen-exposed and control nests (Fig. 3A). We did not include entrance-entrance distance, since our model did not consider surface interactions. The absolute value of the partial least square regression coefficients is proportional to the relative influence of the effect on the response, and signs indicate whether a property is positively or negatively associated with transmission.

## Acknowledgements

We thank Liz Martin-Silverstone and the XTM Facility, Palaeobiology Research Group, University of Bristol, for use and training for micro-CT scanning. We thank Prof. Nicolai Vitt Meyling from the University of Copenhagen for providing the M. brunneum strain. We also thank Andrea Perna for advice on how to extract networks from 3D image data and Rachael Brown, Florent Masson, Thomas Richardson, Daniel Schlppi and Adriano Wanderlingh for comments on the manuscript.

## Contributions

L.L. and N.S designed the study. M.S.A. and K.B. designed and performed the video analysis. M.S.A., K.B. and L.L. conducted the experiment. L.L. designed the code for the network extraction, agent-based model, analysis and figure generation, with input from N.S.. L.L. and N.S. wrote the manuscript.

## Materials availability

Data will be made available here: https://doi.org/10. 5281/zenodo.10278011 All code will be made available here: https://github.com/ Luke-Leckie/Architectural_Immunity_Nest_Networks

## SUPPLEMENTARY MATERIALS

### Supplementary tables and figures to accompany main text

**Figure S1.**
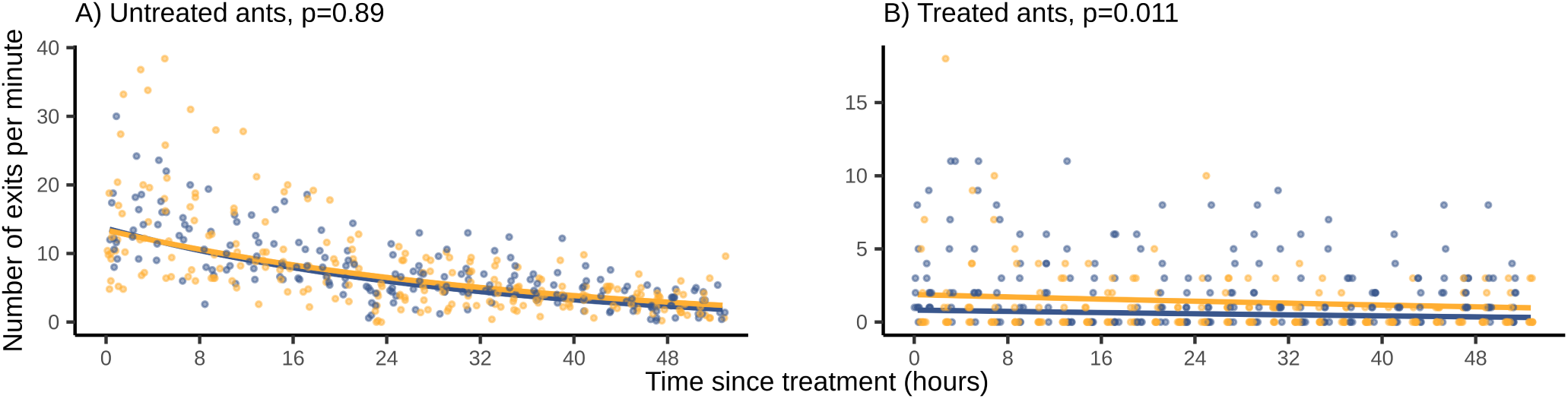
Exit rate of ants. Number of untreated (**A**) and treated (**B**) ants exiting the initial nest-entrance per minute in control (blue) and pathogen-exposed (orange) nests. Each point represents a mean exit rate extracted from a five-minute video (n=446) and lines represent the back-transformed fits. Untreated and treated exits were both log-transformed for statistical analysis. Observations were taken each hour for the first two hours and every two hours thereafter. The significance of the main effect ‘treatment’ in a linear mixed-effects model (LMM) analysis is shown above each graph. Treated workers exited the nest at a significantly higher rate in pathogen-exposed groups. LMM results are provided in Table S1.

**Figure S2.**
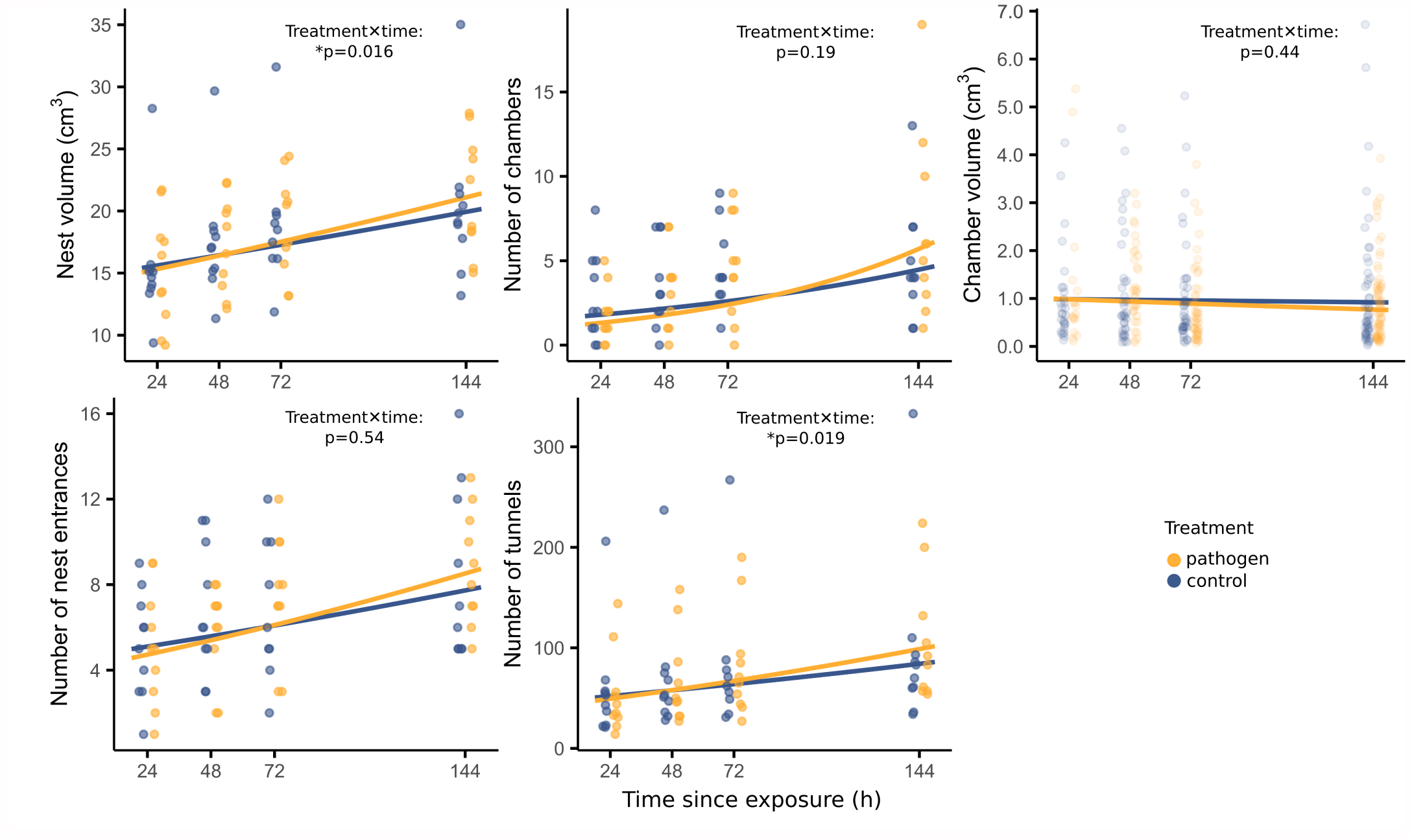
Nest architectural properties as a function of time. Growth and composition of control (blue, left hand points) and pathogen-exposed (orange, right hand points) nests, taken from micro CT nest scans and extracted nest networks. Points represent individual data points (nest properties: n=79, chamber volume: n=343) and lines represent the back-transformed linear model fits. Significance of the interaction between treatment and time is indicated over each graph (*: p<0.05). Nest volume, number of tunnels, and nest entrance number were square-root transformed number of chambers, and chamber volume were log-transformed for statistical analyses. Model coefficients and exact p-values are provided in Table S1.

Table S1. **Results of statistical analyses performed over the post-exposure period.** Significance is highlighted by the colour of the p-value’s cell: no colour p>0.05, yellow p<0.05, blue p<0.01 and green p<0.001. See attached Table S1.

Table S2. **Results of statistical analyses performed on pre-treatment (0h) nests to test for any initial differences.** There was no statistically significant difference between future pathogen-exposed and future control nests for any of the dependent variables tested, as represented by the absence of colour in the p-value cells. See attached Table S2.

**Table S3.**
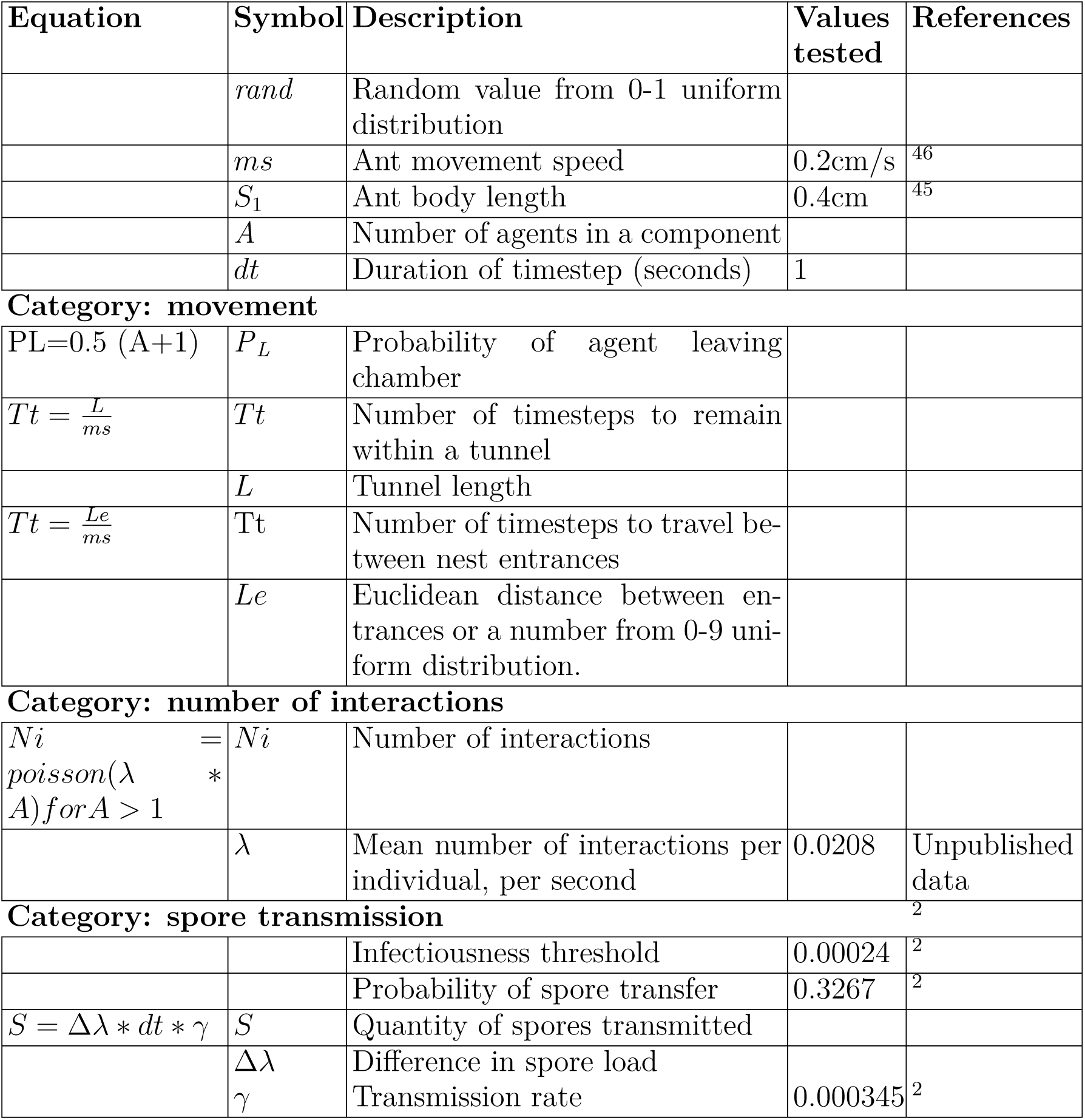
Agent-based model parameters, symbols, and source references.

| Equation | Symbol | Description | Values tested | References |
| --- | --- | --- | --- | --- |
| | $rand$ | Random value from 0-1 uniform distribution | | |
| | $ms$ | Ant movement speed | 0.2cm/s | <sup>46</sup> |
| | $S_1$ | Ant body length | 0.4cm | <sup>45</sup> |
| | $A$ | Number of agents in a component | | |
| | $dt$ | Duration of timestep (seconds) | 1 | |
| **Category: movement** |  |  |  |  |
| PL=0.5 (A+1) | $P_L$ | Probability of agent leaving chamber | | |
| $Tt = \frac{L}{ms}$ | $Tt$ | Number of timesteps to remain within a tunnel | | |
| | $L$ | Tunnel length | | |
| $Tt = \frac{Le}{ms}$ | $Tt$ | Number of timesteps to travel between nest entrances | | |
| | $Le$ | Euclidean distance between entrances or a number from 0-9 uniform distribution. | | |
| **Category: number of interactions** |  |  |  |  |
| $Ni = poisson(\lambda A) for A > 1$ | $Ni$ | Number of interactions | | |
| | $\lambda$ | Mean number of interactions per individual, per second | 0.0208 | Unpublished data |
| **Category: spore transmission** |  |  |  |  |
|  |  | Infectiousness threshold | 0.00024 | <sup>2</sup> |
|  |  | Probability of spore transfer | 0.3267 | <sup>2</sup> |
| $S = \Delta\lambda * dt * \gamma$ | $S$ | Quantity of spores transmitted | | |
| | $\Delta\lambda$ | Difference in spore load | | |
| | $\gamma$ | Transmission rate | 0.000345 | <sup>2</sup> |

### Pathogenand sham-exposure

The generalist fungal entomopathogen *Metarhizium brunneum (* strain MA275, KVL03-143) was used for pathogen exposure. *M. brunneum* is a natural pathogen to *L. niger* and its conidiospores (thereafter ‘spores’) are widely found at high densities in natural soil^48^. Spores adhere to the host cuticle upon contact, then germinate on the host cuticle after 24-48-hours and invade the hemocoel, resulting in host death within 3-7 days^39^, Spores were plated from -80°C stock solutions onto Sabouraud dextrose agar at 23°C for up to three weeks^49^. Spores were harvested in 0.05% Triton-X100 and used within ten days of collection. Prior to use, we checked that the spore germination rate was greater than 90%^2,50^. Pathogenand sham-exposure were performed by dipping individuals in a 1×10^9^spores/ml spore suspension (pathogen) or in a 0.05% Triton-X100 solution only (sham) for five seconds with spring tweezers, and then drying them on filter paper^51^.

### Micro-CT analysis

Nest 3D CT scans were obtained at the XTM Facility, Palaeobiology Research Group, University of Bristol using a Nikon XTH 225 ST X-ray tomography scanner. Scanning conditions were 185kV, 708ms exposure, between 150-190mA current, copper 0.5mm filtration, one frame per projection and a voxel (3D pixel) size of approximately 55.2µm (32.4cm distance). Each scan took approximately 37 minutes to complete. Scanning conditions were chosen to minimise radiation exposure as far as possible whilst ensuring high scan resolution. Time-interval micro-CT scanning at similar doses has been previously used in the closely related ant species *Lasius flavus*, and the scanning procedure did not affect digging behaviour^52^.

### Volume processing

We developed our own image analysis pipeline for the processing of CT-scan data (Fig. S3A). Tiff image stacks of nests were downsampled by a factor of three along each axis and reconstructed using the ORS Dragonfly software. For each nest, scans for each successive day of development were aligned. Noise was reduced by applying 3D gaussian smoothing, with a standard deviation of 1.5. Tunnels were segmented from the substrate by thresholding images at a maximum grayscale intensity value of 16525. The area below the level of the substrate surface was retained for analysis. Samples were processed to remove noise by eroding the tunnels, removing islands under 10,000 voxels in size, re-dilating tunnels, and applying a spherical smoothing operation. Erosion and dilation operations were performed with a kernel size of seven. Smoothing was performed with a kernel size of nine. Tunnel volume and interpolated surface area were obtained from the software directly for each day.

### Network extraction

Volumes were skeletonised in FIJI ImageJ^53^ and further processed using a customwritten Python script. Our network creation pipeline followed the same principles as previously established for social insect nests in Perna *et al. (* 2008)^54^. Initially, skeletons were converted into edge lists where each edge (tunnel) had a true length, representing the distance an ant would have to travel through that edge, and a diameter, measured along, and averaged between, two axes orthogonal to the tunnel midpoint. It was next necessary to post-process these rough networks (Fig. S3B) to more accurately represent the true nest topology. First, we removed all small, spurious edges terminating in a single node and measuring less than 1.4cm in length (“pruning”, Fig. S3B-C). Next, nodes at bends in tunnels were removed by converting any edges measuring less than 0.4cm in length (õne ant body length) into nodes with coordinates at the euclidean midpoint of the edge, and then removing all degree-two non-chamber nodes (Fig. S3C-D; see below for chamber identification method). When edges were converted to nodes or nodes were removed, the new, combined, edge inherited the summed total length of the combined edges and their average diameter. Optimal thresholds for edge concatenation and pruning were determined by selecting parameters which produced networks most closely corresponded to manual network annotations of ten pseudo-randomly selected volumes (two for each scanned time point). Automatic node annotations matched 83.86±11.21% of manual node annotations.

### Node identification

We identified nodes as nest entrances, junctions, endpoints and chambers using a semi-automatic pipeline. First, all chambers were identified using the procedure detailed below. Second, non-chamber nodes whose elevation was within 0.5cm of the highest node in the nest and which were not directly connected to a node of higher elevation were labelled as nest entrances. Any false positive or negative entrance identifications were identified from our scans and corrected manually. Finally, amongst the non-nest entrance nodes, nodes with a degree greater than one were labelled as junctions and nodes with a degree of one were labelled as endpoints.

Chamber identification was performed according to the following semi-automated procedure (Fig. S4). First, nodes which were enlarged compared to the surrounding area and exhibited horizontal floors were identified as candidate chambers^42^. This process was done automatically using the volume thickness maps of 3D images in which filled voxels represented empty space (i.e. the below-ground nest network), with intensity values increasing with the thickness of the area, and empty voxels represented the surrounding substrate (Fig. S3A). The geometric centre of a focal node was located by counting the number of filled voxels within the node along each of its x, y, and z axes (Fig. S4A) and finding the midpoint of each axis iteratively until the diameter no longer increased (Fig. S4B). Measurements of relative enlargement and floor gradient were next taken around that geometric centre. To measure relative enlargement, we created an ellipsoid matching the cavities x/y/z dimensions (‘chamber ellipsoid’) and an ellipsoidal with dimensions 1.75 as large as those of the chamber ellipsoid (‘surroundings ellipsoid’). The intensity values of voxels within the chamber ellpisoid and within the strip separating the chamber ellipsoid and surroundings ellipsoid (‘surroundings shell’) were then sampled, averaged and compared (Fig. S4C) by calculating the intensity ratio *R* = *Ci / Si*, where *Ci* is the mean voxel intensity in the chamber ellipsoid and *Si* is the mean voxel intensity in the surroundings shell, and which represents the relative local enlargement of the focal node. To test whether the focal node had a horizontal floor, the gradient of the cavity floor was calculated along the x and y axes between pairs of points equidistant from the geometric centre. For each pair of points, the number of filled voxels beneath each point was counted to calculate their elevation from the floor. The difference in elevation between the two points was divided by the distance between them, providing a gradient, *g*. This was repeated for all pairs of points along the x and y axes and averaged for each axis to obtain x and y gradients for each prospective chamber (Fig. S4D). The focal node was retained as a candidate chamber if it fulfilled one of the following relative enlargement and gradient conditions for both x and y axes: (1: R>1.3 and *g*<5 percent; 2: R>1.25 and *g*<4 percent; 3: R>1.2 and *g*<3 percent; 4: node mean diameter was more than twice greater than the mean tunnel diameter in the nest and *g*<4 percent). After automatic identification, candidate chamber nodes were labelled as confirmed chambers if the corresponding location on the nest 3D image had been manually annotated as a chamber by blinded observers (Fig. S4E). These criteria were selected from a range of tested combinations of relative thickness/gradient thresholds as resulting in chamber assignments most closely corresponding to manual annotations by blinded observers. Over the 100 nests networks, 90.50% of manual annotations also satisfied algorithmic criteria and were defined as chambers.

We tracked the development of nest networks over time by linking node identities between pairs of networks corresponding to successive stages of nest development (i.e., between networks at 0h and 24h, 24h and 48h, 48h and 72h and 72h and 144h postexposure). This was accomplished by assigning unique identifications to each node in the earlier nest network and matching it with the nearest node in the new network, within a maximum distance of 1.4cm (as was used for edge pruning). Nodes in the new network were assigned new identities if the closest node in the earlier network was more than 1.4cm away, or if it could be paired to a closer node in the new network. We further improved node tracking over time by applying the following rules: end nodes in the new network could only be matched to end nodes in the earlier network, junctions in the new network could only be matched to junctions or end nodes in the earlier network, and chambers in the new network could be matched to end nodes, chambers, or junctions in the earlier network.

**Figure S3.**
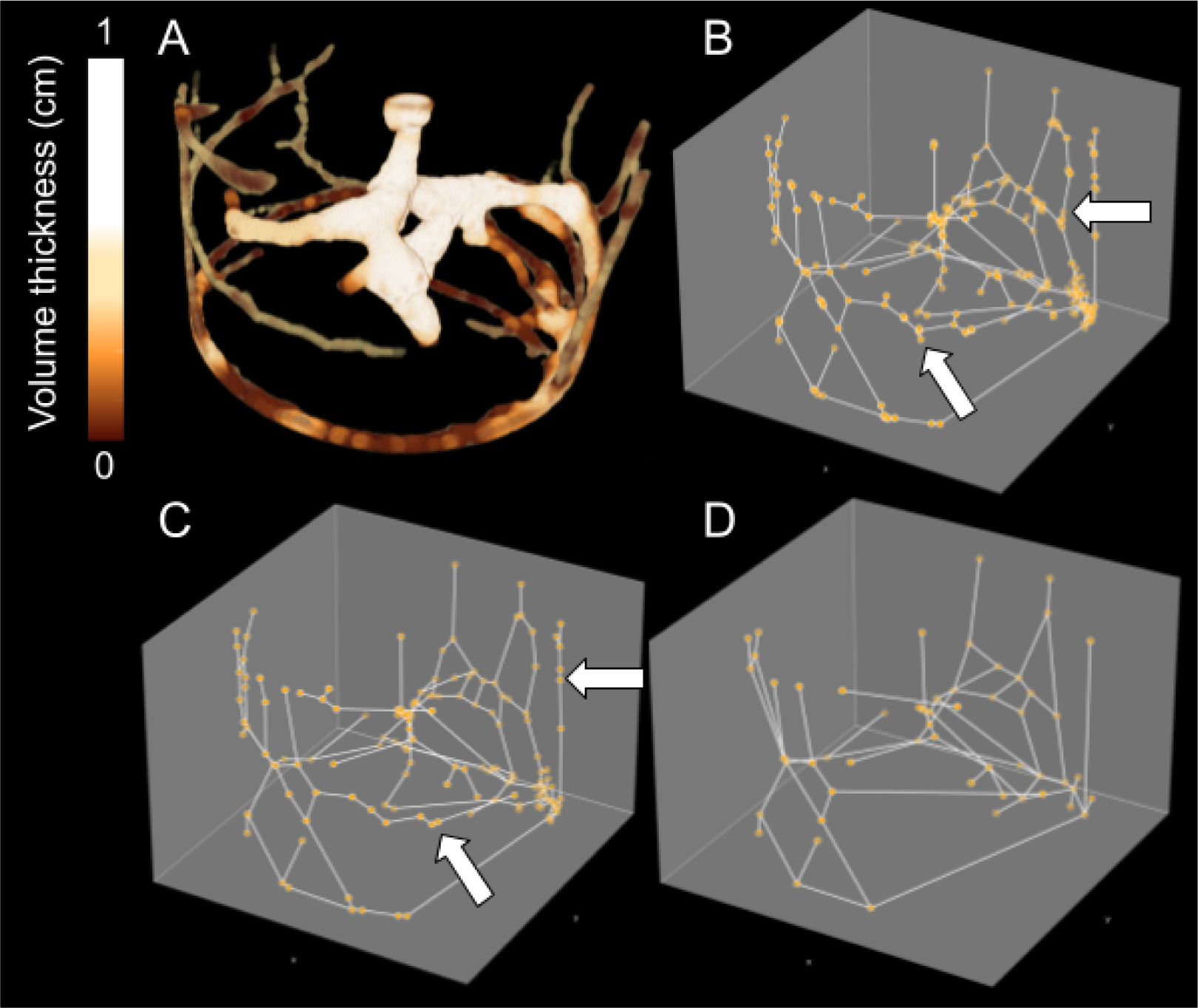
Network extraction procedure. **A:** Volume thickness map of nest micro CT-scan, with brighter regions indicating increased thickness; **B:** network of skeletonised volume. Junctions, dead-ends, entrances, and chambers are represented in orange. Network has spurious branches, indicated by arrows; **C:** pruned network, with endpoint edges <1.4cm long removed. Network still has nodes at tunnel bends, indicated by arrows; **D:** final network with degree-two nodes removed.

### Agent-based model pseudocode

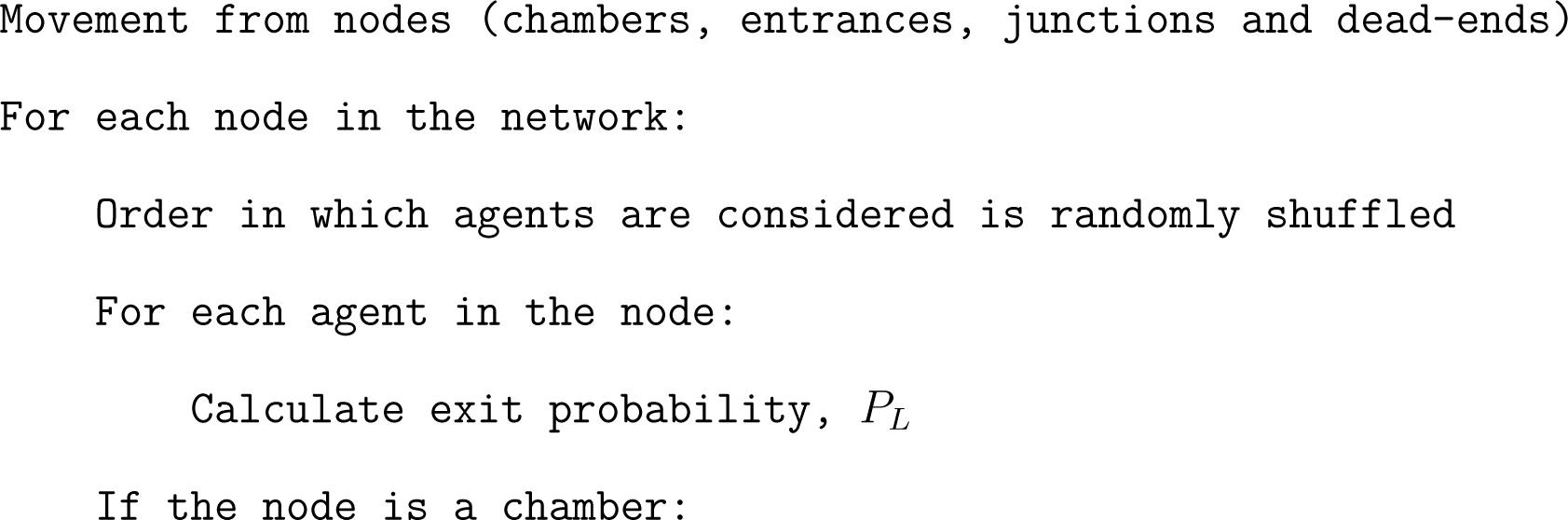

**Figure S4.**
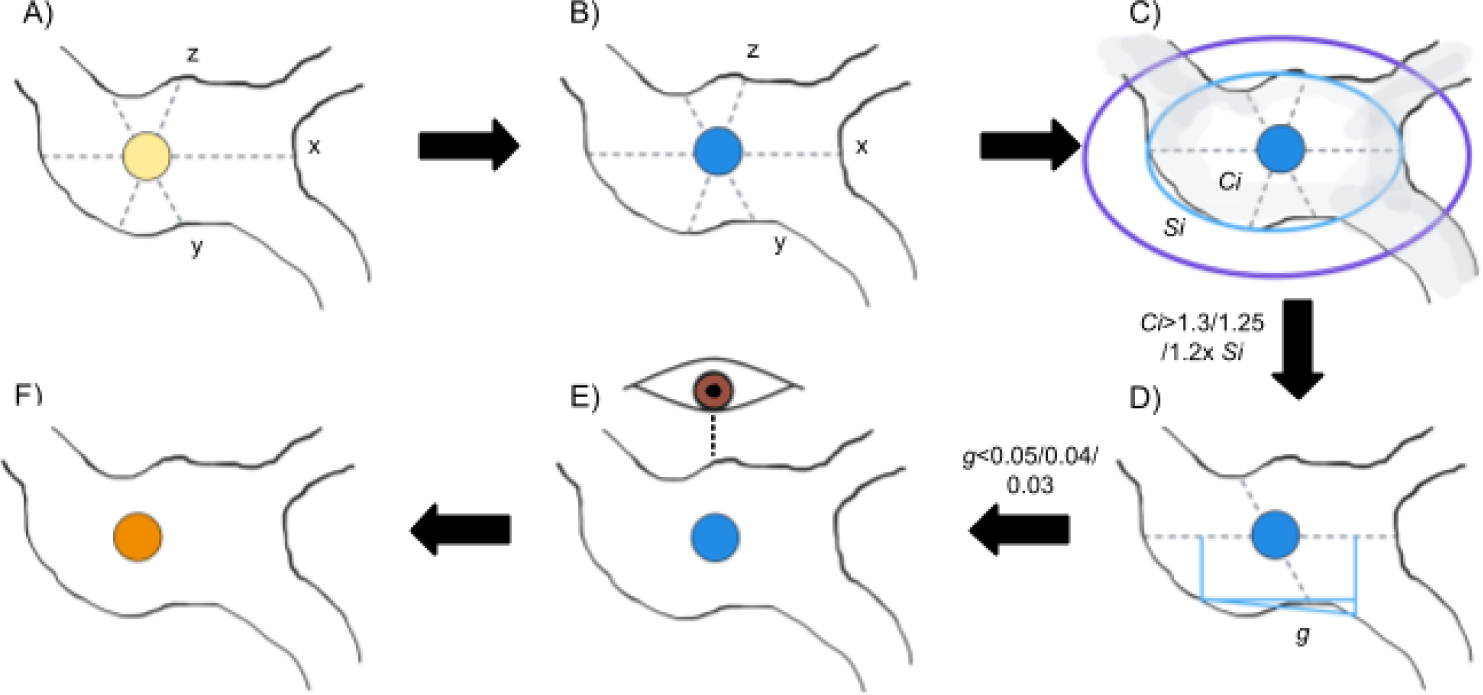
Flowchart of chamber identification pipeline. **A:** For a focal node (yellow circle) within a cavity (bounded by black lines), x, y, and z axes are calculated (dotted lines); **B:** the central point of the focal node, at which the x, y, and z radii are symmetrical, is located (geometric centre, blue circle); **C:** we calculated the mean voxel intensity *Ci* within the chamber ellipsoid (cyan ellipse) and the mean voxel intensity *Si* within the surrounding shell (between cyan and purple ellipses), whose diameter was 1.75x the size of the chamber ellipsoid. Wider regions of the nest are more white and have higher voxel values; **D:** the gradient, *g (* diagonal blue line), was calculated between equidistant pairs of points along x and y axes. If *Ci* is >1.3x Si and *g*<5 percent, *Ci* is >1.25x *Si* and *g*<4 percent, *Ci i*s >1.2x *Si* and *g*<3 percent or focal node diameter more than 2x mean nest tunnel diameter and g<4 percent, the focal node was identified as candidate chamber; **E:** if the algorithmically identified candidate chamber annotation was confirmed by a manual annotation by a human observer (eye) the node was labelled as a chamber (**E**, orange circle).

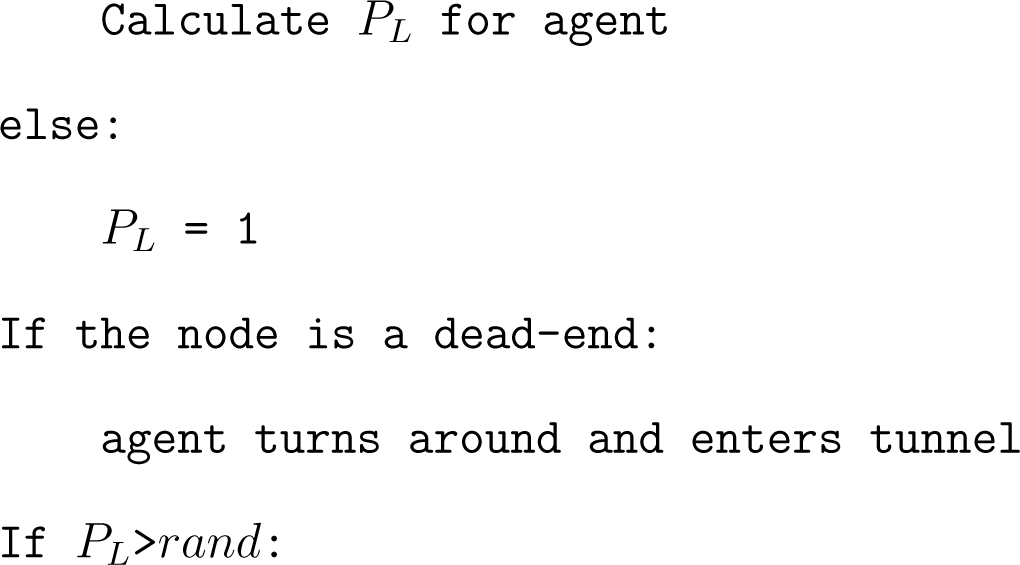

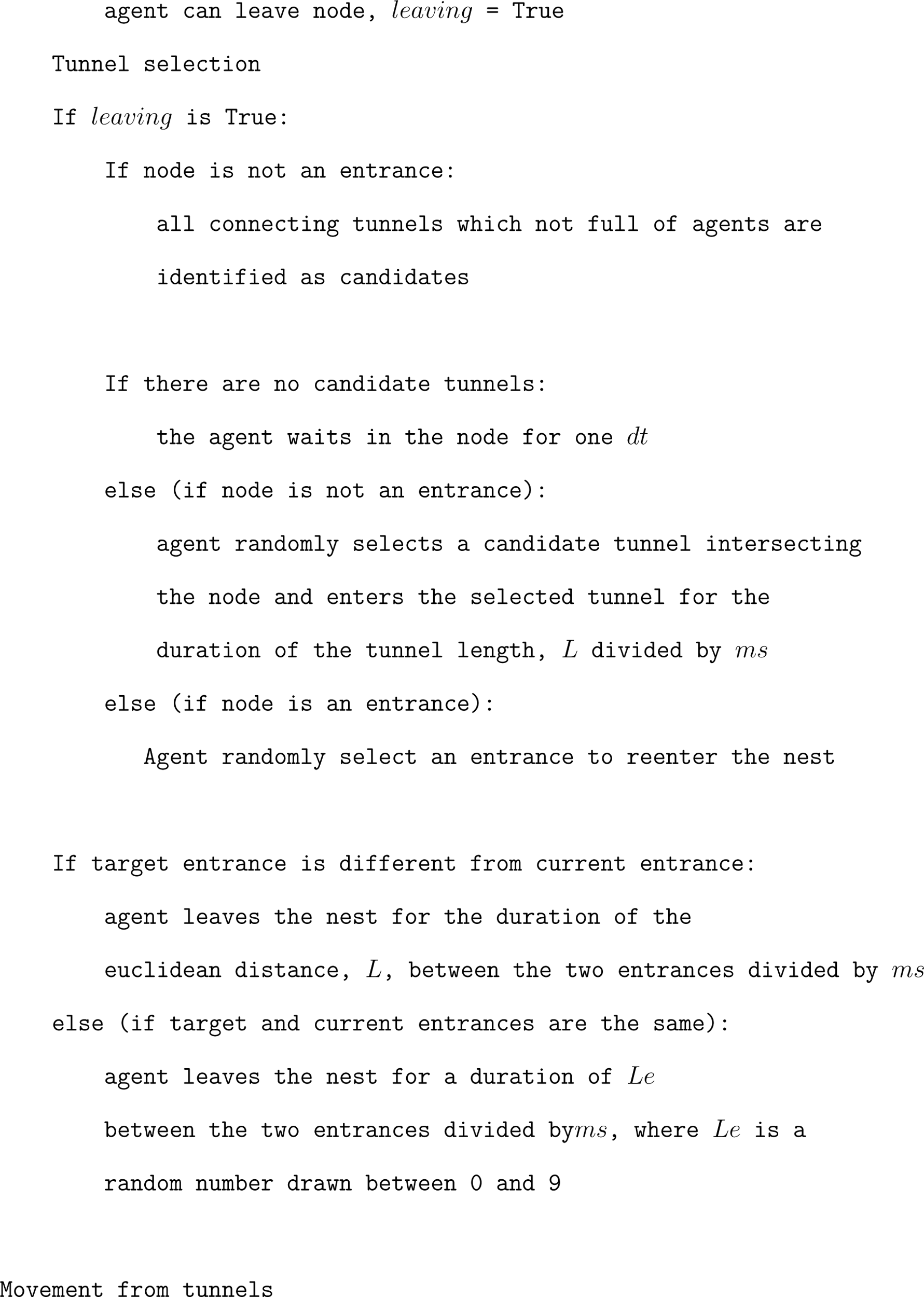

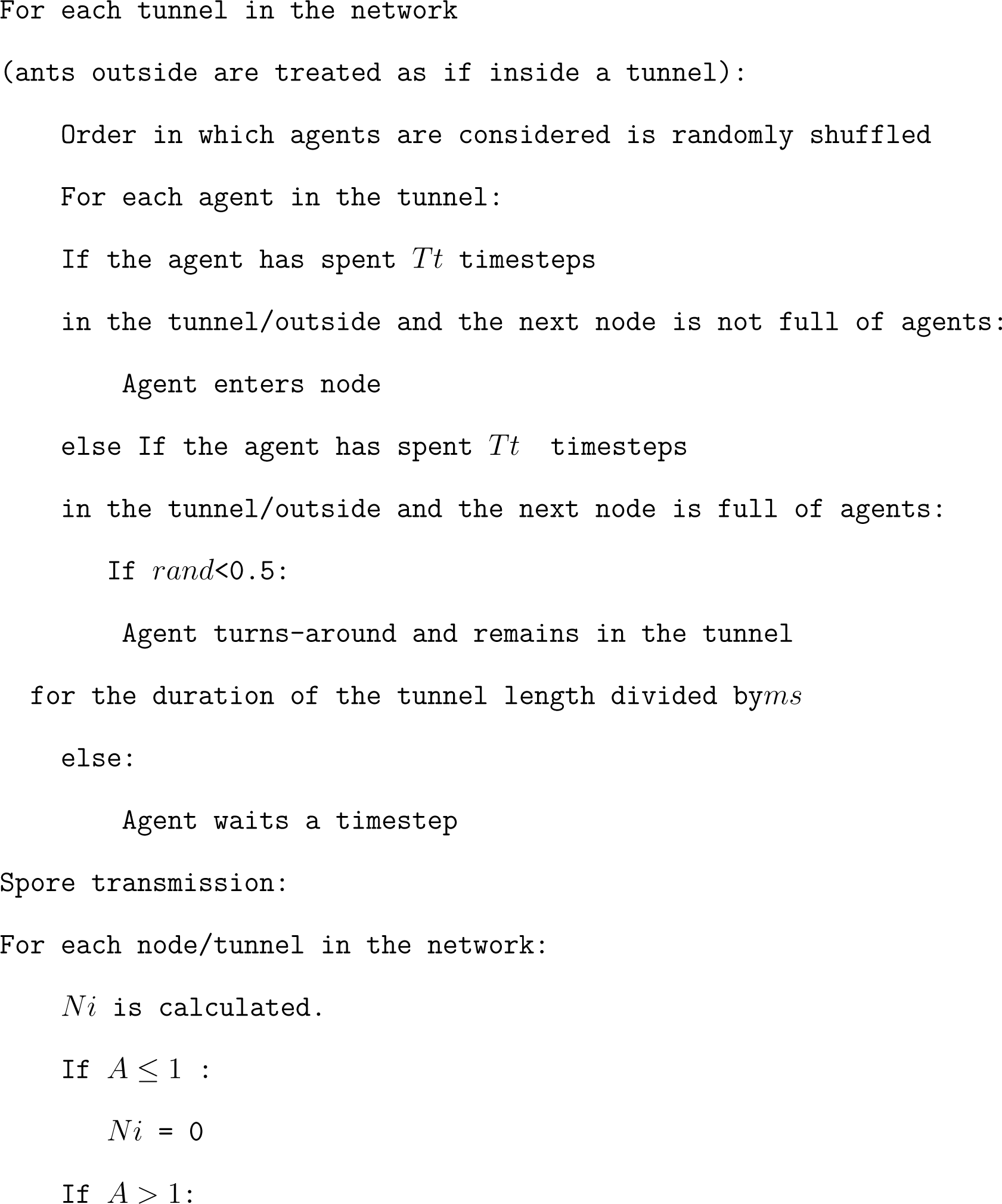

## REFERENCES

1. Danon, L., House, T. A., Read, J. M. & Keeling, M. J. Social encounter networks: collective properties and disease transmission. Journal of The Royal Society Interface 9, 2826–2833 (2012).

2. Stroeymeyt, N. et al. Social network plasticity decreases disease transmission in a eusocial insect. Science 362, 941–945. https://www.science.org/doi/full/10.1126/science.aat4793 (2024) (Nov. 2018).

3. De Freslon, I., Martínez-López, B., Belkhiria, J., Strappini, A. & Monti, G. Use of social network analysis to improve the understanding of social behaviour in dairy cattle and its impact on disease transmission. Applied animal behaviour science 213, 47–54 (2019).

4. Tracey, J. A., Bevins, S. N., VandeWoude, S. & Crooks, K. R. An agent-based movement model to assess the impact of landscape fragmentation on disease transmission. en. Ecosphere 5, art119. issn: 2150-8925. https://onlinelibrary.wiley.com/doi/abs/ 10.1890/ES13-00376.1 (2024) (2014).

5. Silva, G. C. & Ribeiro, E. M. S. The impact of Brazil’s transport network on the spread of COVID-19. en. Scientific Reports 13, 2240. issn: 2045-2322. https://www.nature.com/articles/s41598-022-27139-1 (2024) (Feb. 2023).

6. Uddin, S., Khan, A., Lu, H., Zhou, F. & Karim, S. Suburban Road Networks to Explore COVID-19 Vulnerability and Severity. en. International Journal of Environmental Research and Public Health 19, 2039. issn: 1660-4601. https://www.mdpi.com/1660-4601/19/4/2039 (2024) (Jan. 2022).

7. Keeling, M. J., Danon, L., Vernon, M. C. & House, T. A. Individual identity and movement networks for disease metapopulations. Proceedings of the National Academy of Sciences 107, 8866–8870. https://www.pnas.org/doi/abs/10.1073/pnas.1000416107(2024) (May 2010).

8. Colizza, V., Barrat, A., Barthélemy, M. & Vespignani, A. The role of the airline transportation network in the prediction and predictability of global epidemics. Proceedings of the National Academy of Sciences 103, 2015–2020. https://www.pnas.org/doi/ 10.1073/pnas.0510525103 (2024) (Feb. 2006).

9. Stephenson, J. F., Perkins, S. E. & Cable, J. Transmission risk predicts avoidance of infected conspecifics in Trinidadian guppies. Journal of Animal Ecology 87, 1525–1533 (2018).

10. Lopes, P. C., Block, P. & König, B. Infection-induced behavioural changes reduce connectivity and the potential for disease spread in wild mice contact networks. Scientific reports 6, 31790 (2016).

11. Bos, N., Lefèvre, T., Jensen, A. & d’Ettorre, P. Sick ants become unsociable. Journal of evolutionary biology 25, 342–351 (2012).

12. Heiler, G., et al. *Country-wide mobility changes observed using mobile phone data during COVID-19 pandemic* in 2020 *IEEE international conference on big data (big data)* (IEEE, Dec. 2020), 3123–3132. isbn: 978-1-72816-251-5. https://ieeexplore.ieee.org/document/9378374/ (2023).

13. Fisher, T. Viral cities. Places Journal. issn: 21647798. https://placesjournal.org/article/viral-cities/ (2024) (Oct. 2010).

14. Crucitti, P., Latora, V. & Porta, S. Centrality measures in spatial networks of urban streets. Physical Review. E, Statistical, Nonlinear, and Soft Matter Physics 73, 036125. 10.1103/%7BPhysRevE%7D.73.036125(2023) (Mar. 2006).

15. Megahed, N. A. & Ghoneim, E. M. Antivirus-built environment: Lessons learned from Covid-19 pandemic. Sustainable Cities and Society 61, 102350. issn: 22106707. https://linkinghub.elsevier.com/retrieve/pii/S2210670720305710 (2023) (Oct. 2020).

16. Cremer, S., Armitage, S. A. O. & Schmid-Hempel, P. Social Immunity. English. Current Biology 17, R693–R702. issn: 0960-9822. https://www.cell.com/current-biology/abstract/S0960-9822(07)01503-5 (2024) (Aug. 2007).

17. Perna, A. & Theraulaz, G. When social behaviour is moulded in clay: on growth and form of social insect nests. Journal of Experimental Biology 220, 83–91. 10.1242/jeb.143347 (2023) (Jan. 2017).

18. Stroeymeyt, N., Casillas-Pérez, B. & Cremer, S. Organisational immunity in social insects. Current Opinion in Insect Science 5, 1–15 (2014).

19. Leclerc, J.-B., Pinto Silva, J. & Detrain, C. Impact of soil contamination on the growth and shape of ant nests. Royal Society open science 5, 180267 (2018).

20. Kim, M. et al. Lessons from a COVID-19 hospital, republic of korea. Bulletin of the World Health Organization 98, 842–848. 10.2471/%7BBLT%7D.20.261016 (2023) (Dec. 2020).

21. Salathé, M. & Jones, J. H. Dynamics and Control of Diseases in Networks with Community Structure. en. PLOS Computational Biology 6, e1000736. issn: 1553-7358. https://journals.plos.org/ploscompbiol/article?id=10.1371/journal.pcbi.1000736 (2024) (Apr. 2010).

22. Latora, V. & Marchiori, M. Efficient behavior of small-world networks. Physical Review Letters 87, 198701. 10.1103/%7BPhysRevLett%7D.87.198701 (2023) (Nov. 2001).

23. Barthélemy, M., Barrat, A., Pastor-Satorras, R. & Vespignani, A. Dynamical patterns of epidemic outbreaks in complex heterogeneous networks. Journal of Theoretical Biology 235, 275–288. 10.1016/j.jtbi.2005.01.011 (2023) (July 2005).

24. Kiss, I. Z., Green, D. M. & Kao, R. R. The network of sheep movements within Great Britain: Network properties and their implications for infectious disease spread. *Journal of the Royal Society*, Interface 3, 669–677. 10.1098/rsif.2006.0129 (2023) (Oct. 2006).

25. Sah, P., Leu, S. T., Cross, P. C., Hudson, P. J. & Bansal, S. Unraveling the disease consequences and mechanisms of modular structure in animal social networks. Proceedings of the National Academy of Sciences 114, 4165–4170. https://www.pnas.org/doi/abs/10.1073/pnas.1613616114 (2024) (Apr. 2017).

26. Volz, E. M., Miller, J. C., Galvani, A. & Meyers, L. A. Effects of Heterogeneous and Clustered Contact Patterns on Infectious Disease Dynamics. en. PLOS Computational Biology 7, e1002042. issn: 1553-7358. https://journals.plos.org/ploscompbiol/article?id=10.1371/journal.pcbi.1002042 (2024) (June 2011).

27. Newman, M. J. A measure of betweenness centrality based on random walks. Social networks 27, 39–54. issn: 03788733. http://linkinghub.elsevier.com/retrieve/pii/S0378873304000681 (2017) (Jan. 2005).

28. Wang, S., Du, Y. & Deng, Y. A new measure of identifying influential nodes: Efficiency centrality. Communications in Nonlinear Science and Numerical Simulation 47, 151–163. issn: 10075704. http://linkinghub.elsevier.com/retrieve/pii/S1007570416304129 (2023) (June 2017).

29. Mouronte-López, M. L. Analysing the vulnerability of public transport networks. Journal of Advanced Transportation 2021, 1–22. issn: 2042-3195. https://www.hindawi.com/journals/jat/2021/5513311/ (2023) (Mar. 2021).

30. Dudkina, E. et al. A comparison of centrality measures and their role in controlling the spread in epidemic networks. International journal of control 97, 1325–1340. issn: 0020-7179. https://www.tandfonline.com/doi/full/10.1080/00207179.2023.2204969 (2024) (May 2023).

31. Kissler, S. M. et al. Geographic transmission hubs of the 2009 influenza pandemic in the United States. Epidemics 26, 86–94. 10.1016/j.epidem.2018.10.002 (2024) (Mar. 2019).

32. Gravish, N., Gold, G., Zangwill, A., Goodisman, M. A. & Goldman, D. I. Glass-like dynamics in confined and congested ant traffic. Soft matter 11, 6552–6561 (2015).

33. Chang, J., Powell, S., Robinson, E. J. H. & Donaldson-Matasci, M. C. Nest choice in arboreal ants is an emergent consequence of network creation under spatial constraints. en. Swarm Intelligence 15, 7–30. issn: 1935-3820. 10.1007/s11721-021-00187-5 (2024) (June 2021).

34. Stockmaier, S., et al. Infectious diseases and social distancing in nature. Science 371. issn: 0036-8075. https://www.sciencemag.org/lookup/doi/101126/science.abc8881 (2023) (Mar. 2021).

35. Cook, Z., Franks, D. W. & Robinson, E. J. H. Efficiency and robustness of ant colony transportation networks. Behavioral Ecology and Sociobiology 68, 509–517. issn: 0340-5443. http://link.springer.com/10.1007/s00265-013-1665-8 (2023) (Mar. 2014).

36. Pie, M. R., Rosengaus, R. B. & Traniello, J. F. A. Nest architecture, activity pattern, worker density and the dynamics of disease transmission in social insects. Journal of Theoretical Biology 226, 45–51. issn: 0022-5193. https://www.sciencedirect.com/science/article/pii/S0022519303003096 (2024) (Jan. 2004).

37. Ulrich, Y., Saragosti, J., Tokita, C. K., Tarnita, C. E. & Kronauer, D. J. C. Fitness benefits and emergent division of labour at the onset of group living. Nature 560, 635– 638. issn: 0028-0836. http://www.nature.com/articles/s41586-018-0422-6 (2023) (Aug. 2018).

38. Sloane, N. J. A. & Plouffe, S. The encyclopedia of integer sequences https://cir.nii.ac.jp/crid/1130000794892856704 (2024) (Academic Press, 1995).

39. Grizanova, E. V., Coates, C. J., Dubovskiy, I. M. & Butt, T. M. Metarhizium brunneum infection dynamics differ at the cuticle interface of susceptible and tolerant morphs of Galleria mellonella. Virulence 10, 999–1012. issn: 2150-5594. 10.1080/21505594.2019.1693230 (2024) (Jan. 2019).

40. Liu, J., Liew, S. S., Wang, J. & Pu, K. Bioinspired and biomimetic delivery platforms for cancer vaccines. Advanced Materials 34, e2103790. 10.1002/adma.202103790 (2023) (Jan. 2022).

41. Toffin, E., Kindekens, J. & Deneubourg, J.-L. Excavated substrate modulates growth instability during nest building in ants. Proceedings. Biological Sciences / the Royal Society 277, 2617–2625. 10.1098/rspb.2010.0176 (2023) (Sept. 2010).

42. Pinter-Wollman, N. Nest architecture shapes the collective behaviour of harvester ants. Biology Letters 11, 20150695. https://royalsocietypublishing.org/doi/full/10.1098/rsbl.2015.0695 (2024) (Oct. 2015).

43. Hagberg, A., Swart, P. J. & Schult, D. A. *Exploring network structure, dynamics, and function using NetworkX* tech. rep. (Los Alamos National Laboratory (LANL), Los Alamos, NM (United States), 2008).

44. Clauset, A., Newman, M. E. & Moore, C. Finding community structure in very large networks. *Physical Review E—Statistical*, Nonlinear, and Soft Matter Physics 70, 066111 (2004).

45. Khuong, A. et al. Stigmergic construction and topochemical information shape ant nest architecture. Proceedings of the National Academy of Sciences 113, 1303–1308 (2016).

46. Richardson, T. O., Stroeymeyt, N., Crespi, A. & Keller, L. Two simple movement mechanisms for spatial division of labour in social insects. en. Nature Communications 13, 6985. issn: 2041-1723. https://www.nature.com/articles/s41467-022-34706-7 (2024) (Nov. 2022).

47. Cohen, J. *Statistical Power Analysis for the Behavioral Sciences* 2nd ed. isbn: 978-0-203-77158-7 (Routledge, New York, July 1988).

48. Steinwender, B. M., Enkerli, J., Widmer, F., Eilenberg Jand Thorup-Kristensen, K. & Meyling, N. V. Molecular diversity of the entomopathogenic fungal Metarhizium community within an agroecosystem. Journal of Invertebrate Pathology 123, 6–12. 10.1016/j.jip.2014.09.002 (2023) (Nov. 2014).

49. Walker, T. N. & Hughes, W. O. H. Adaptive social immunity in leaf-cutting ants. Biology Letters 5, 446–448. 10.1098/rsbl.2009.0107 (2023) (Aug. 2009).

50. Konrad, M. et al. Ants avoid superinfections by performing risk-adjusted sanitary care. Proceedings of the National Academy of Sciences 115, 2782–2787. https://www.pnas.org/doi/abs/10.1073/pnas.1713501115 (2024) (Mar. 2018).

51. Alciatore, G. et al. Immune challenges increase network centrality in a queenless ant. Proceedings. Biological Sciences / the Royal Society 288, 20211456. 10.1098/rspb.2021.1456 (2023) (Sept. 2021).

52. Minter, N. J., Franks, N. R. & Robson Brown, K. A. Morphogenesis of an extended phenotype: four-dimensional ant nest architecture. Journal of The Royal Society Interface 9, 586–595. https://royalsocietypublishing.org/doi/abs/10.1098/rsif.2011.0377 (2024) (Aug. 2011).

53. Schindelin, J., et al. Fiji: an open-source platform for biological-image analysis. Nature Methods 9, 676–682. 10.1038/nmeth.2019 (2021) (June 2012).

54. Perna, A. et al. Topological efficiency in three-dimensional gallery networks of termite nests. Physica A: Statistical Mechanics and its Applications 387, 6235–6244. issn: 03784371. http://linkinghub.elsevier.com/retrieve/pii/S0378437108006675 (2023) (Oct. 2008).

